# Cerebral Venous Blood Flow Regulates Brain Fluid Clearance via Dural Lymphatics

**DOI:** 10.1101/2024.12.12.628127

**Authors:** Marie-Renee El Kamouh, Myriam Spajer, Ruchith Singhabahu, Kurt A Sailor, Marie-Charlotte Bourrienne, Laura Mouton, Sunil Koundal, Diana Doukhi, Abel Grine, Justus Ninneman, Amelle Nasry, Anne-Laure Joly Marolany, David Akbar, Joshua Gottschalk, Anthony Ruze, Elora Buscher, Dominique Langui, Jerome Van Wassenhove, Mikael Mazighi, Anne Eichmann, Pierre-Marie Lledo, Helene Benveniste, Mathieu Santin, Stéphane Lehericy, Jean-Léon Thomas, Stéphanie Lenck

## Abstract

The vascular system regulates brain clearance through arterial blood flow and lymphatic drainage of cerebrospinal fluid (CSF). Idiopathic intracranial hypertension (IIH), characterized by elevated intracranial pressure and dural venous sinus stenoses, can be treated by restoring venous blood flow via venous stenting, suggesting a role for venous blood flow in brain fluid clearance. Using magnetic resonance imaging (MRI) in IIH patients and healthy controls, we identified that dural venous stenoses in IIH were associated with impaired lymphatic drainage, perivenous fluid retention, and brain fluid accumulation. To investigate this further, we developed a mouse model with bilateral jugular vein ligation (JVL), which recapitulated key human findings, including intracranial hypertension, calvarial lymphatic regression, and brain swelling due to impaired clearance. To further dissect the respective roles of dural lymphatics and venous blood flow in brain clearance, we performed JVL in mice with dural lymphatic depletion. These mice exhibited spontaneous elevated intracranial pressure, but JVL did not further exacerbate this effect. Moreover, the synchronous restoration of brain clearance and dural lymphatics observed in mice after JVL was absent in lymphatic-deficient mice.Transcriptomic analyses revealed that lymphatic remodeling induced by JVL was driven by VEGF-C signaling between dural mesenchymal and lymphatic endothelial cells. These findings establish the dural venous sinuses as a critical platform where venous blood flow interacts with mesenchymal cells to preserve dural lymphatic integrity and function, essential for brain fluid clearance.

## Introduction

Idiopathic intracranial hypertension (IIH) is a neurological disorder characterized by headaches and elevated intracranial pressure (ICP) in the absence of a brain tumor or other identifiable disease. ^1^ IIH predominantly affects young women, often in the context of weight gain or obesity, suggesting a role for hormonal factors in its pathogenesis. Patients with IIH exhibit bilateral stenoses of the dural transverse venous sinuses, resulting in impaired cerebral venous outflow. Venous stenting effectively relieves venous obstruction, normalizes ICP, and alleviates symptoms, indicating that disrupted venous blood flow contributes to abnormal brain fluid drainage in these patients. ^2^ However, the mechanisms by which cerebral venous outflow influences brain fluid clearance remain poorly understood.

The defective clearance of brain waste contributes to toxic protein accumulation and neurodegenerative diseases. ^3^ Because the brain is devoid of intrinsic lymphatic vasculature, brain clearance requires a special waste-removal system. A mechanistic model has been proposed that combines intracerebral CSF drainage by the glymphatic system with extracerebral drainage of glymphatic outflow by lymphatic vessels. ^4^ The glymphatic system implicates the CSF as the mediator of brain waste clearance, circulating within the perivascular spaces and the brain parenchyma under the regulatory control of perivascular astrocytes. ^5,6^ The lymphatic circuit that drains the intracerebral glymphatic outflow includes the dural and extracranial vessels and their collecting cervical lymph nodes. ^4,7–10^ CSF entry into the periarterial spaces is controlled, at least in part, by arterial pulsation. ^11,12^ The outflow pathways of intracerebral CSF, their regulatory mechanisms, and the specific role of perivenous spaces in the context of brain clearance remain controversial topics. ^13^

Based on this premise, we hypothesized that abnormal blood flow in the dural venous sinuses may alter the properties of perisinusal spaces in IIH patients, and that such alterations may extend to extracerebral lymphatic vessels that regulate the clearance of CNS-derived fluids from the brain. By using contrast-enhanced MRI in IIH patients and healthy controls, along with a mouse model of altered cerebral venous outflow via bilateral jugular vein ligation (JVL), we confirmed that brain clearance depends on dural lymphatics and demonstrated that cerebral venous outflow is essential for maintaining the integrity and function of these vessels. These results provide critical insights into IIH pathophysiology and establish a biological foundation for understanding the interplay between cerebral venous and lymphatic systems.

## Results

### Occlusion of dural venous sinuses impairs perivascular fluid drainage in IIH patients

We performed contrast-enhanced 3T-MRI using black blood imaging with the 3D T1 SPACE DANTE ^9^ sequence to detect low-velocity fluid drainage along dural perivascular spaces in IIH patients and healthy volunteers.^9,14–16^ This technique enables the visualization of perisinusal drainage as a proxy measurement of lymphatic drainage, as histological findings support the presence of lymphatic vessels within perisinusal spaces in humans, primates and mice. ^9,14^ This Perisinusal flow imaging combined with other sequences were applied to cohorts of twenty patients with IIH and twenty healthy and age-matched volunteers.

The participants were imaged with a 3T Prisma MRI scanner (Siemens Healthineers) using a series of sequences administered either before, or after, intravenous (IV) injection of the contrast agent, gadobutrol (Fig. 1A). The venous elliptical sequence was used to delineate the dural venous sinuses and the major bridging veins, while the 3D T1 SPACE DANTE sequence was employed to determine the volume of perisinusal and perivenous fluids (Fig. 1B). ^9^ We successfully imaged thirty six of the forty participants, including sixteen IIH patients and nineteen healthy volunteers. After post-processing of the MRI scans, we analyzed the three-dimensional reconstruction of the venous sinuses (blue) and perisinusal compartments (green) (Fig.1 C, D).

**Figure 1.**
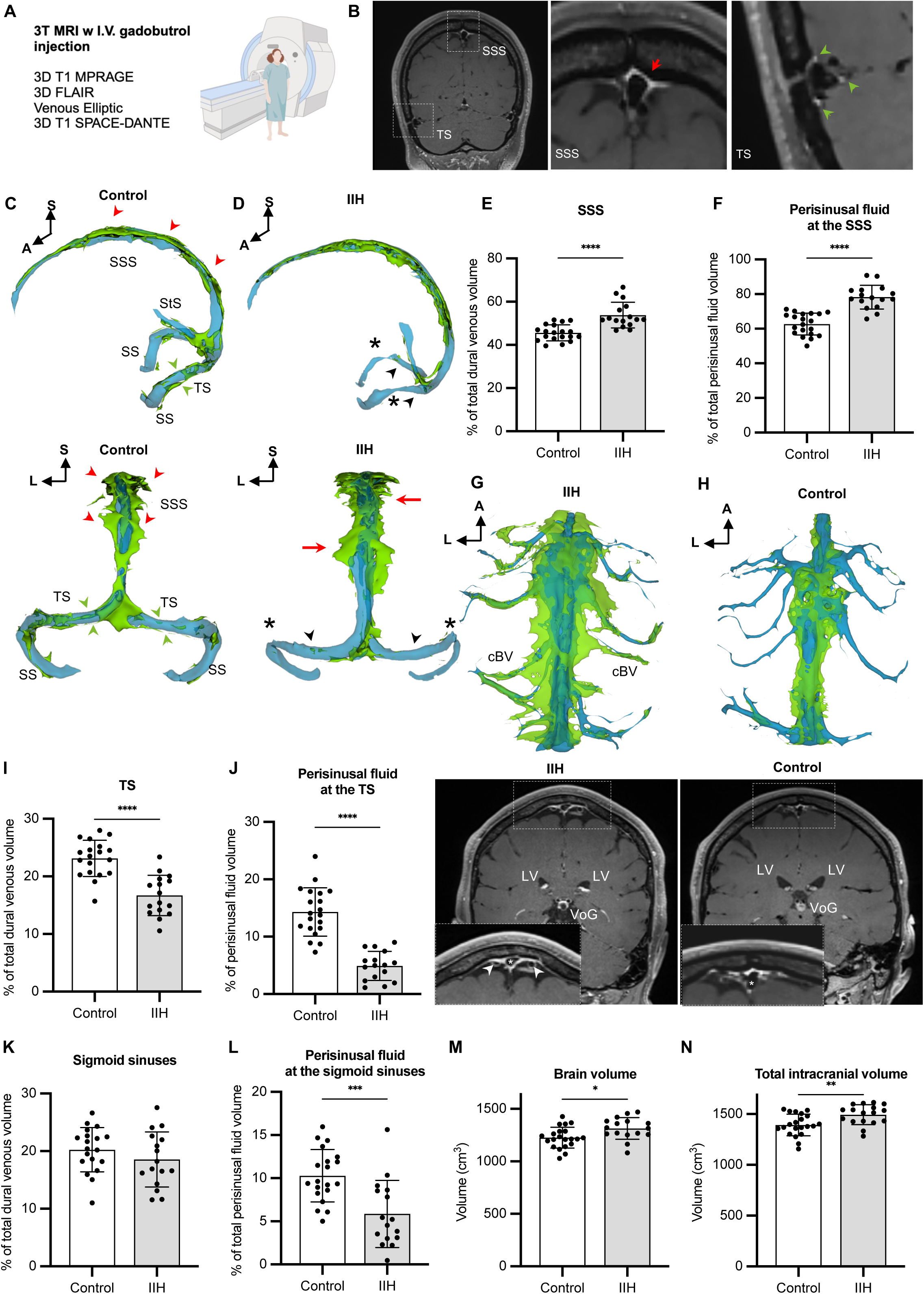
Alterations of dural venous sinus and perisinus associate with brain oedema in IIH patients. (A) 3T MRI protocol in IIH patients and healthy controls includes the following sequences: 3D T1 MPRAGE (magnetization-prepared rapid acquisition gradient-echo); MR venography with a venous elliptic; 3D FLAIR (fluid-attenuated inversion recovery), and 3D T1 SPACE DANTE. (B) Coronal 3D T1 SPACE DANTE scan of the head crossing the superior sagittal sinus (SSS) and the transverse sinus (TS). Global view (left panel). Magnification of the perisinus enhancement showing cisternae at the SSS (center panel, red arrowhead) and tubular-like structures at the TS (right panel, green arrowhead). (C, D) 3D reconstructions of both MR venography (blue) and 3D T1 SPACE DANTE (green) scans in healthy volunteers (C) and IIH patients (D). Lateral (top panels) and dorsal (bottom panels) views. Dural tubular-like structures, interpreted as lymphatic vessels (C, green arrowheads), disappear in IIH patients (D, black arrowhead), notably at the level of venous stenoses (D, blue asterisk). Conversely, perisinusal cisternae at the SSS (red arrowheads) are significantly enlarged in IIH patients (D) compared to controls (C). (E, F) Quantification showing increased volumes of SSS (E) and perisinus (F) in IIH patients. (G, H) Paravenous signal enhancement along cortical bridging veins (cBVs) in IIH patients (G) compared to healthy controls (H). Top panels: 3D reconstruction of merged MR venography (blue) and 3D T1 SPACE DANTE (green). IIH patients show continuous signal enhancement between the enlarged perisinus cisternae of the dura and the paravenous spaces along cBVs in the subarachnoid space (I, white arrowhead), albeit not healthy volunteers (J). Bottom panels: native 3D T1 SPACE DANTE coronal scans at the level of the SSS and perisinus (dotted frame, magnified in the insert (white asterisk)). Vein of Galen (VoG) and lateral ventricles (LV). (I-L) Volume quantification at the level of the TS (I, J) and sigmoid sinus (SS) (K, L). Venous volume (I, K). Perisinus volume (J, L). In IIH patients, perisinus volume (J, L) is reduced, together with venous stenosis in the TS (I). (M, N) The brain (M) and intracranial (N) volumes are increased in IIH patients. Unpaired t-test in E, F and I to N. \**P* < 0.05, \*\**P* < 0.01, \*\*\**P* < 0.001, \*\*\*\**P* < 0.0001. Error bars indicate SD. Elements generated with: BioRender (https://biorender.com) (A); 3D-Slicer (https://www.slicer.org) (B, C, D, G, H).

The venous sinuses and perisinusal spaces had similar volumes in both IIH patients and healthy controls (SF. 1A, B), despite the presence of bilateral obstructions at the TS in all IIH patients (Fig.1 D, asterisk). Controls showed perisinusal cisternae (red arrowheads in Fig. 1B, C) along the superior sagittal sinus (SSS), as well as vessel-like tubular structures along the transverse (TS), sigmoid (SS), and straight sinuses (green arrowheads, Fig. 1B, C). In contrast, IIH consistently displayed two different local alterations of the dural sinus system. First, IIH patients exhibited enlarged SSS and perisinusal cisternae (red arrowheads), as compared to healthy controls (Fig. 1D, E and F). Additionally, perivenous enhancement along the cortical bridging veins located in the subarachnoid space was detected in patients (white arrowheads, Fig. 1G) but not visible in healthy volunteers (Fig. 1H). Therefore, the superior sagittal perisinus is enlarged and continuous with the cortical perivenous spaces in IIH patients. Second, downstream venous sinuses and perisinusal spaces in the latero-posterior region of the dural sinus system were both reduced in IIH patients as compared to controls (Fig. 1C-D, I-L). The bilateral constriction of transverse sinuses (as denoted by asterisks) was consistently associated with the lack of perisinusal drainage in IIH patients (Fig. 1D, black arrowheads). Therefore, obstruction of the transverse sinus in IIH patients impaired drainage of perivascular spaces along the cerebral bridging veins and the dural sagittal perisinus. These observations show that perivascular fluid is continuous between cerebral bridging veins and dural sinuses and depends upon cerebral venous outflow.

We confirmed that intracranial hypertension was linked to altered brain fluid homeostasis in IIH patients through volumetric analyses. Using the 3D-FLAIR sequence, we monitored the volumes of the brain, ventricles, and choroid plexuses, while the total intracranial volume was extracted from the 3D T1 MPRAGE sequence. Our findings revealed that brain volume was increased in IIH patients compared to controls (Fig. 1M), even though ventricular volume remained similar between the two groups (SF. 1C). Interestingly, IIH patients also exhibited larger intracranial volumes than controls (Fig. 1N). This may be attributed to brain swelling and could result from the thinning of cranial bone previously reported in IIH patients.^17^

Altogether these observations validate the hypothesis that, in IIH patients, alteration of the dural venous blood flow is associated with abnormal perivenous drainage. The restriction of perisinusal fluid drainage in the transverse sinuses logically correlates with upstream retention of fluids in perisinusal spaces along the dural superior sagittal sinus and in brain tissues. Intracranial hypertension may be the logical outcome of this significant brain fluid retention that was measured in patients with IIH.

### Surgical ligation of venous outflow in mice causes cerebral oedema and impairment of brain fluid clearance

Based on our findings in IIH patients, we speculated that alterations in venous sinus outflow could impair function of the adjacent dural lymphatic, potentially compromising brain fluid drainage. To investigate this, we developed a surgical model of cerebral venous outflow obstruction in mice.

Adult female mice underwent bilateral ligation of the internal and external jugular veins (JVL mice) (Fig. 2A). The jugular veins are responsible for draining the cerebral blood toward the subclavian veins, while the cervical paravertebral venous plexuses serve as the only alternative route. We first conducted MRI monitoring of venous remodeling using a 11.7T MRI scanner at various time points after JVL surgery (2, 7, 14, 28, and 60 days) and after sham surgery (Fig. 2B). 2D Time-of-flight (TOF) MR venography revealed significant remodeling of the facial veins located upstream of the ligation points compared to sham operated mice (Fig. 2C). The diameter of these extracranial veins doubled within 2 days post-surgery (2-dps) and gradually returned to baseline over the course of 4 weeks (Fig. 2D). The volume of the dural venous sinuses remained unchanged throughout the same period (Fig. 2E).

**Figure 2.**
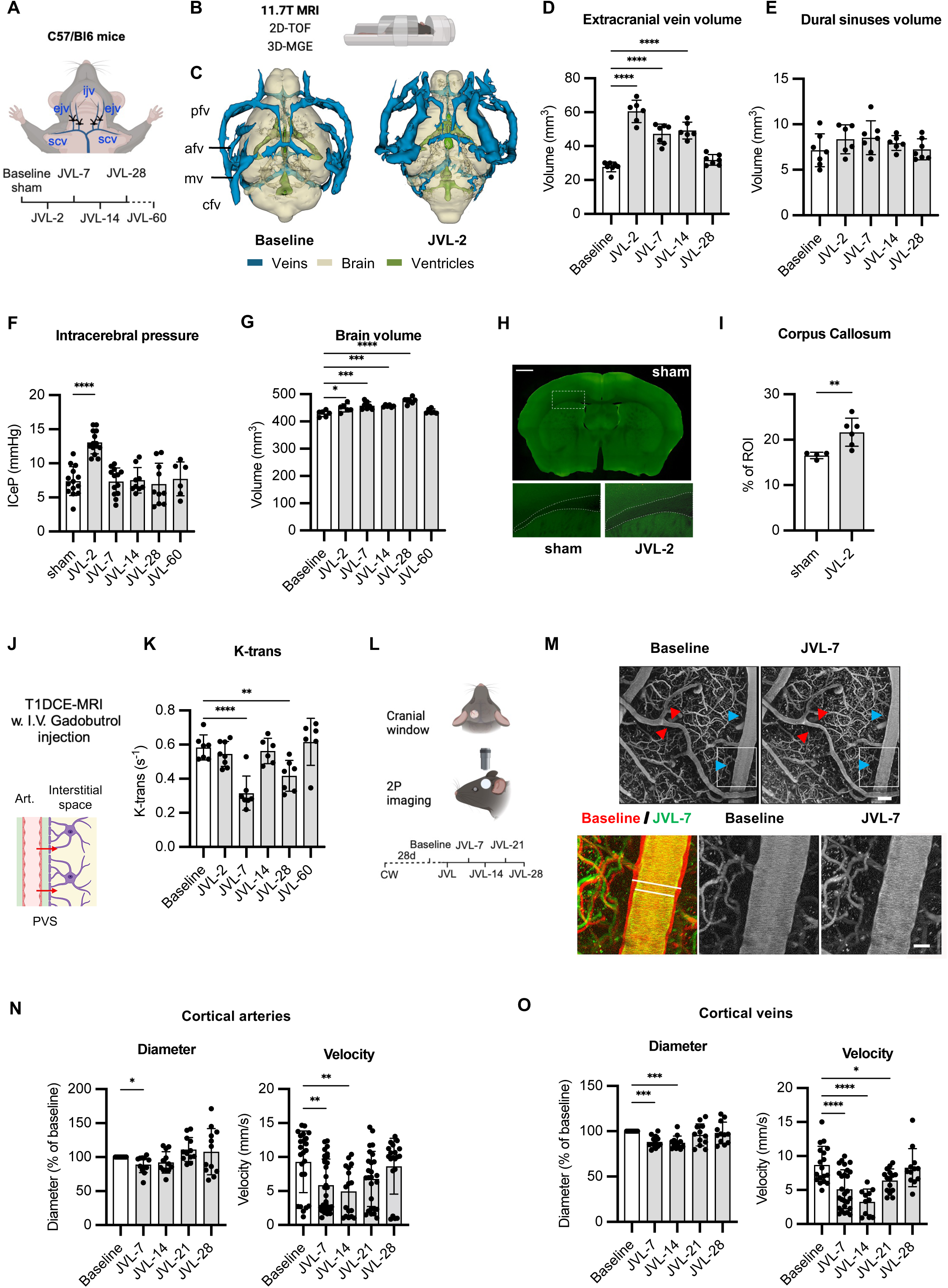
Jugular vein ligation induces a transient fluctuation in cerebral blood flow and brain oedema in mice. (A) Adult female mice underwent blockade of cerebral venous outflow by jugular vein ligation (JVL mice). The external (ejv) and internal jugular veins (ijv) are ligated prior to their confluence in the subclavian vein (scv). JVL mice assessment at different endpoints (2-, 7-, 14-, 28-, 60-dps). (B) The following sequences are performed on a 11.7 T MRI: 3D time-of-flight (TOF) for venous system reconstruction; 3D multi-gradient echo (MGE) sequence for brain and ventricle volume reconstruction. (C-E) 3D reconstruction and quantification of extracerebral and dural veins. Imaging of intra- and extracranial veins (blue), brain (grey), and ventricles (green) in a baseline mouse and a mouse at 2 days post-surgery (dps), highlighting the enlargement of extracranial veins (posterior facial vein [pfv], anterior facial vein [afv], maxillary vein [mv] joining the common facial vein [cfv]) (C). Enlargement of extracranial veins begins at 2 dps and gradually normalizes by 28 dps (D). Dural venous sinus volume remains stable over time (E). (F) JVL induces an acute and transient increase in intracerebral pressure (ICeP) at 2-dps, followed by normalization. (G) JVL induces a long-lasting brain swelling initiated at 2-dps, peaking at 28-dps, and returning to baseline by 60-dps. (H,I) Cerebral oedema correlates with enlargement of corpus callosum (H) assessed by surface area measurements by contrast on auto-fluorescent brain slices (I). (J,K) Assessment of BBB integrity by K-trans measurement. T1 dynamic contrast-enhanced (DCE)-MRI was performed on a 11.7 T scanner after IV. injection of gadobutrol (J). K-trans measures the passage of gadobutrol from arteries to the cerebral interstitial (extravascular and extracellular) space (J). K-trans value does not increase after jugular ligation, ruling out BBB leakage. K-trans decreases at 7- and 14-dps, indicating that transfer of intravascular water into the cerebral interstitial space is reduced (K). (L-O) Imaging of cortical vasculature using two-photon microscopy in mice with jugular ligation. A cranial window (CW) was placed over the somatosensory/motor cortex 28 days prior to JVL, and imaging was performed at baseline, 7-, 14-, 21- and 28-dps after IV. injection of fluorescent dextrans (n = 2 mice) (F). (G) Two-photon images illustrating vasoconstriction in both arteries (red arrowheads) and veins (blue arrowheads). Higher magnification of a cortical vein demonstrates reduced venous diameter in JVL-7 mice. (H, I) Quantification of vascular diameter and velocity in veins (H) and arteries (I). Note vasoconstriction and decreased velocity in arteries at 7- and 14-dps, while velocity in veins was decreased from 7- to 21-dps. Ordinary one-way ANOVA with Dunnett’s multiple comparisons test was used in D to G, K, N (velocity), O (velocity). Repeated measures ANOVA with Dunnett’s multiple comparisons test was used in N and O for vessel diameters. Mann Whitney U-test was used in I. \**P* < 0.05, \*\**P* < 0.01, \*\*\**P* < 0.001, \*\*\*\**P* < 0.0001. Error bars indicate SD. Scale bars: 50 µm (M, top), 20 µm (M, bottom) and 800 µm (H). Elements in A, B, J and L were created using BioRender (https://biorender.com). Elements in C were generated using 3D-Slicer (https://www.slicer.org).

To determine whether impaired cerebral venous outflow altered brain fluid homeostasis, we measured intracerebral pressure (ICeP) over time, using a pressure sensor catheter inserted into the brain parenchyma in isoflurane anesthetized mice. A significant, yet transient, increase of ICeP was observed at 2-dps, followed by a rapid return to baseline at 7-dps (Fig. 2F). To assess wether this elevation of ICeP correlated with an increase of brain volume, as observed in humans (Fig. 1M), we measured brain volume with 11.7 Tesla MRI. 3D magnetization-prepared gradient echo (MGE) sequence revealed that brain volume progressively increased from the time of surgery until 28-dps, and returned to baseline only at 60-dps (Fig. 2M). Brain volume expanded without changing the ventricular volume during the same period (SF. 2A). Hence, the blockage of cerebral venous outflow causes cerebral edema that persists beyond the initial phase of intracranial hypertension. We determined that, at the peak of ICeP, brain swelling affected the axonal tracts rather than the grey matter, as was observed by surface area measurements of the corpus callosum contrast on auto-fluorescent brain slices at the level of the somatosensory cortex in sham operated and JVL-2 mice (Fig. 2H, I). To conclude, bilateral jugular vein ligated mice develop a transient IIH-like phenotype characterized by transient intracranial hypertension and persistent brain edema predominantly in white matter tracts.

To investigate whether leakage of the blood brain barrier (BBB) might contribute to the cerebral edema observed after JVL, the BBB integrity of brain sections was assessed postmortem after IV. administration of dextran fluorescent tracers with two different molecular weights (3 and 40 kDa). Dextran did not diffuse from the blood into surrounding tissues, or around dural venous sinuses at 2-dps after JVL (SF. 2B, C), suggesting that the BBB and the dural sinuses maintain their integrity after JVL. We also examined the transverse sinuses of JVL-7 mice by electron microscopy and observed no venous endothelium break, in agreement with the finding that small size blood-injected tracer did not leak around the sinus in JVL-2 mice (SF. 2D, E). We also assessed the BBB in vivo over time using dynamic contrast-enhanced (DCE)-MRI after IV injection of gadobutrol. A T1-weighted perfusion sequence (DCE-FLASH) was used to measure the volume transfer constant (K-Trans) which served as an indicator of BBB permeability (Fig. 2J). K-Trans quantifies gadolinium transport between plasma and intracerebral tissues, encompassing both extracellular and extravascular spaces. ^18^ Following JVL, there was no increase in K-Trans, ruling out excessive gadobutrol entry into brain tissue and indicating an intact BBB. Instead, K-Trans decreased at 7- and 28-dps (Fig. 2K), reflecting reduced water transport from blood to brain. This decrease in vascular permeability, despite pressure and fluid buildup in brain tissue, suggests that brain oedema following JVL results from impaired fluid clearance and potential disruption of CSF drainage through cerebral tissue.

To explore the observed decrease in microvascular permeability detected by DCE-MRI, we examined the brain vasculature using two-photon microscopy (Fig. 2L-O) and light-sheet fluorescence microscopy (LSFM) (SF. 2F, G). Two-photon imaging was conducted through a cranial window placed over the motor/somatosensory cortex, enabling sequential imaging of cortical vessels from baseline to 28-dps. At each imaging time point, fluorescent dextran was injected once into the retro-orbital vein to highlight the vascular network (Fig. 2L). As shown in Fig. 2M-O, cortical arteries slightly constricted at 7-dps (Fig. 2N), while veins constricted at 7- and 14-dps (Fig. 2O), with both returning to baseline later. Blood flow velocity in arteries and veins decreased after JVL and normalized by 21-dps for arteries and 28-dps for veins (Fig. 2N, O). This reflects a transient reduction in blood perfusion in the fronto-parietal cortex draining into the SSS. Additionally, we used LSFM imaging of whole brains to obtain a broader view of the brain arterial vasculature labeled with antibodies against smooth muscle actin (SMA) to highlight perivascular smooth muscle cells that are enriched around arteries. Compared to sham operated mice (SF. 2F), JVL-14 mice showed more cortical arterio-arterial anastomoses at the cortical surface (stars), between the middle cerebral and anterior cerebral arteries, demonstrating local remodeling of cerebral arterial circuits upon JVL (SF. 2G). We speculate these arterio-arterial anastomoses may ensure redistribution of blood supply to prevent hypoxia in the frontal cortical regions.

### Rerouting of lymphatic CSF drainage after JVL

Having established a surgical IIH model in mice, we examined if JVL induced remodeling of the dural lymphatic circuitry by whole-mount imaging of the dural lymphatic vasculature in skullcap and skull-base samples (Fig. 3A). A transgenic Prox1-Cre-tdTomato reporter mouse^19^ was used to detect Prox1 expressing-cells (Fig. 3B), while anti-Lyve-1 antibodies were used to stain samples isolated from C57Bl/6J mice, with or without JVL (Fig. 3C). The lymphatics of the dorsal and baso-ventral areas of the skull both responded synchronously but in an opposite manner to JVL. During the first week, the dorsal lymphatic bed regressed in JVL mice compared to controls, with a decrease expression of Prox1 and Lyve-1 at the level of confluence of sinus (COS) (Fig. 3D, E) and smaller size lymphatics in the dorsal part of transverse sinus (Fig. 3F), while no size change was observed along the SSS (Fig. 3G). Conversely, lymphatic vessels were dilated in the ventral part of transverse sinus (Fig. 3H) and the basal part of the skull (Fig. 3 I-K). At two weeks after JVL, the dural lymphatic vasculature was restored in both regions of the skull. The two week-period of reduced cerebral blood perfusion is therefore associated with a global remodeling of dural lymphatic circuits.

**Figure 3.**
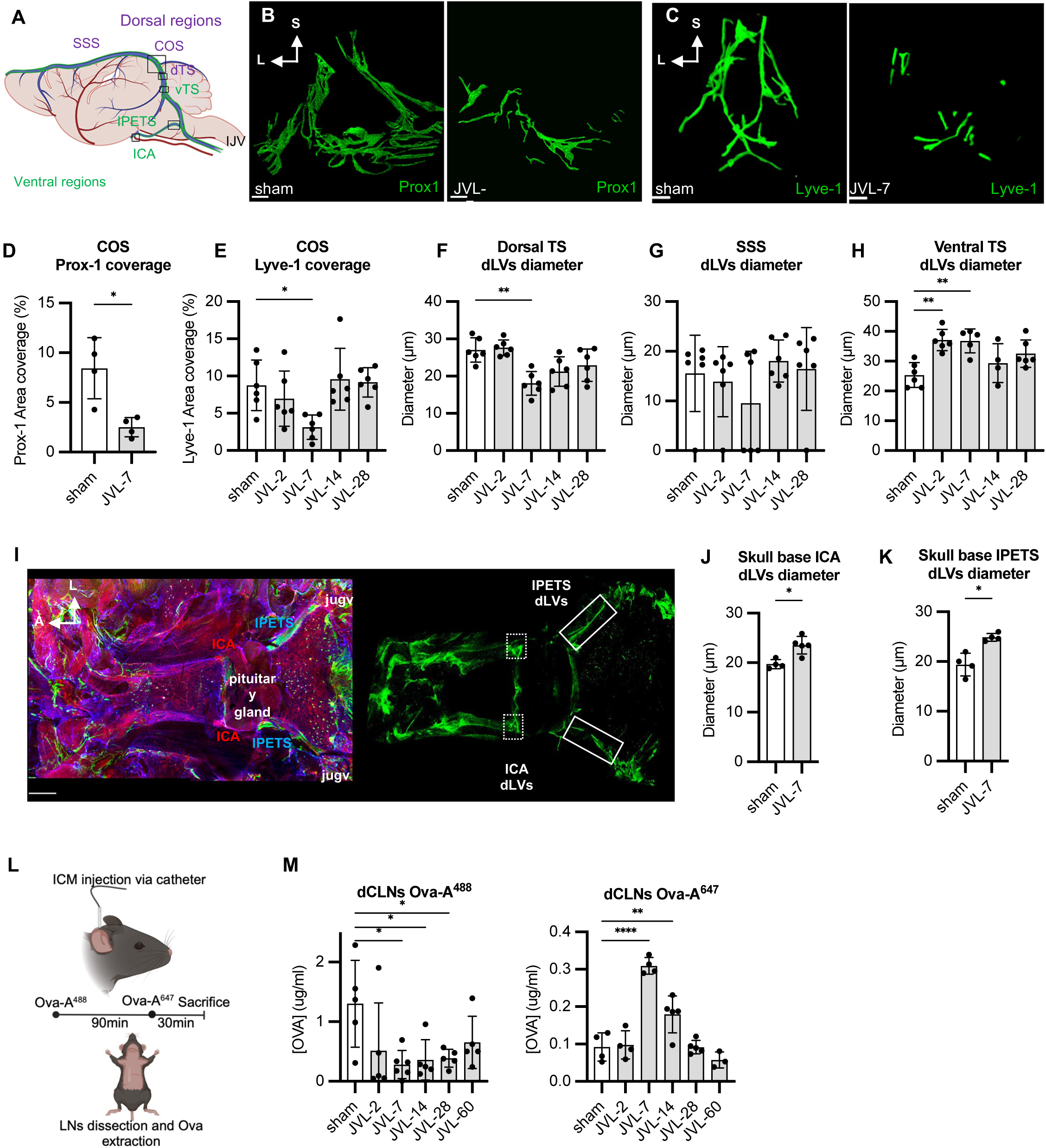
Dural lymphatic remodeling after jugular vein ligation. (A) Dural lymphatics were examined in the calvaria (B-G), i.e. around the superior sagittal sinus (SSS), confluence of sinuses (COS), and dorsal transverse sinus (dTS), as well as in the latero-basal regions of the skull (H-K), i.e. around the ventral transverse sinus (vTS), inferior petrosal sinus (iPETS), and internal carotid arteries (ICA). (B, G). Calvaria lymphatics. Regression of COS lymphatic pattern at 7-dps in JVL mice (B, C). Prox1-dTomato mice (B) and Lyve-1 immunolabeling (C). (D, E) Quantification of Prox1 and Lyve-1 fluorescence areas shows recovery of calvaria lymphatics at 14-dps. (F) The diameter of dTS lymphatics decreased at 7-dps before returning to baseline at 14-dps. (G) No change in the diameter of SSS lymphatics after JVL. (H, K) Latero-basal dural lymphatics. The diameter of vTS lymphatics increased at 2- and 7-dps before normalization at14-dps (H). Confocal imaging of the skull base after immunolabeling with anti-podocalyxin (PDLX) and -Lyve-1 antibodies to identify blood and lymphatic vessels, respectively (I). internal carotid arteries (ICA); inferior petrosal sinuses (iPETS). Dural lymphatic diameter increased at 7-dps around the ICA (J) and along the iPETS (K). (L, M). Kinetics of CSF lymphatic drainage after jugular ligation. CSF drainage into dCLNs was evaluated after ICM. injection of both OVA-A^488^ and OVA-A^647^ at 120 min and 30 min, respectively, before sacrifice (L). JVL mice showed CSF drainage alterations (M). At 7-dps, less OVA-A^488^ marker remained after 120 min, while OVA-A^647^ was increased after 30 min. CSF drainage was normalized by 60-dps. Ordinary one-way ANOVA with Dunnett’s multiple comparisons test was used in E to H, M. Mann Whitney U-test was used in D, J, K. \**P* < 0.05, \*\**P* < 0.01, \*\*\**P* < 0.001, \*\*\*\**P* < 0.0001. Error bars indicate SD. Scale bars: 200 µm (B, C), 500 µm (I). Elements in A and L were created using BioRender (https://biorender.com).

We next measured CSF drainage into the brain-draining lymph nodes after JVL, using intra-cisterna magna (ICM) injection of a fluorescent Alexa-ovalbumin (OVA) marker and subsequent dosage of marker content in collecting lymph nodes, to evaluate the kinetics of CSF drainage. ^20^ JVL and sham operated mice received successive ICM injections of two different markers, a green OVA-A^488^ and a far-red OVA-A^647^ conjugate that were delivered at 120 minutes before sacrifice for OVA-A^488^, and at 30 minutes for OVA-A^647^. OVA-conjugates were measured in several CSF-collecting lymph nodes, including deep and superficial cervical lymph nodes as well as lumbo-aortic and sacral lymph nodes (Fig. 3L). From 7- to 28-dps, we measured a reduction in OVA-A^488^ content 120 minutes after ICM injection, and an increase in OVA-A^647^ content 30 minutes after ICM injection in all collecting lymph nodes, indicating that CSF drainage from the meninges was accelerated upon JVL and dural lymphatic remodeling (Fig. 3M, SF3). After 28-dps, the flow rate gradually decreased in the lymph nodes, normalizing completely by two months post-surgery. These data indicate that the JVL induced dural lymphatic remodeling and accelerated ventral CSF drainage into cervical lymph nodes.

### Synchronized remodeling of dural lymphatic and intracerebral CSF drainage

To investigate which mechanisms may cause the acceleration of extracranial CSF drainage after JVL, we analyzed the intracerebral circulation of CSF at the peak of lymphatic drainage alteration (7-dps). DCE-MRI was performed on JVL and sham operated mice with a 9.4T scanner, using a T1 (T1 DCE-MRI) sequence at one hour after ICM injection of gadoteric acid (Gd-DOTA), as previously reported (Fig. 4A). ^20–22^ Quantitative analysis of the T1 maps showed that the transport of Gd-DOTA into the brain parenchyma decreased in JVL mice compared to sham operated mice (Fig. 4 B, C). The acceleration of CSF drainage into cervical lymph nodes (Fig. 3O) thus correlated with a reduction of intracerebral CSF drainage, as previously reported. ^23^ Reduced CSF entering the brain may cause excessive volume of extracerebral CSF that requires accelerated drainage to cervical lymph nodes. However, in JVL mice, extracranial transport of Gd-DOTA through the cribriform plate was not significantly increased (SF3).

**Figure 4.**
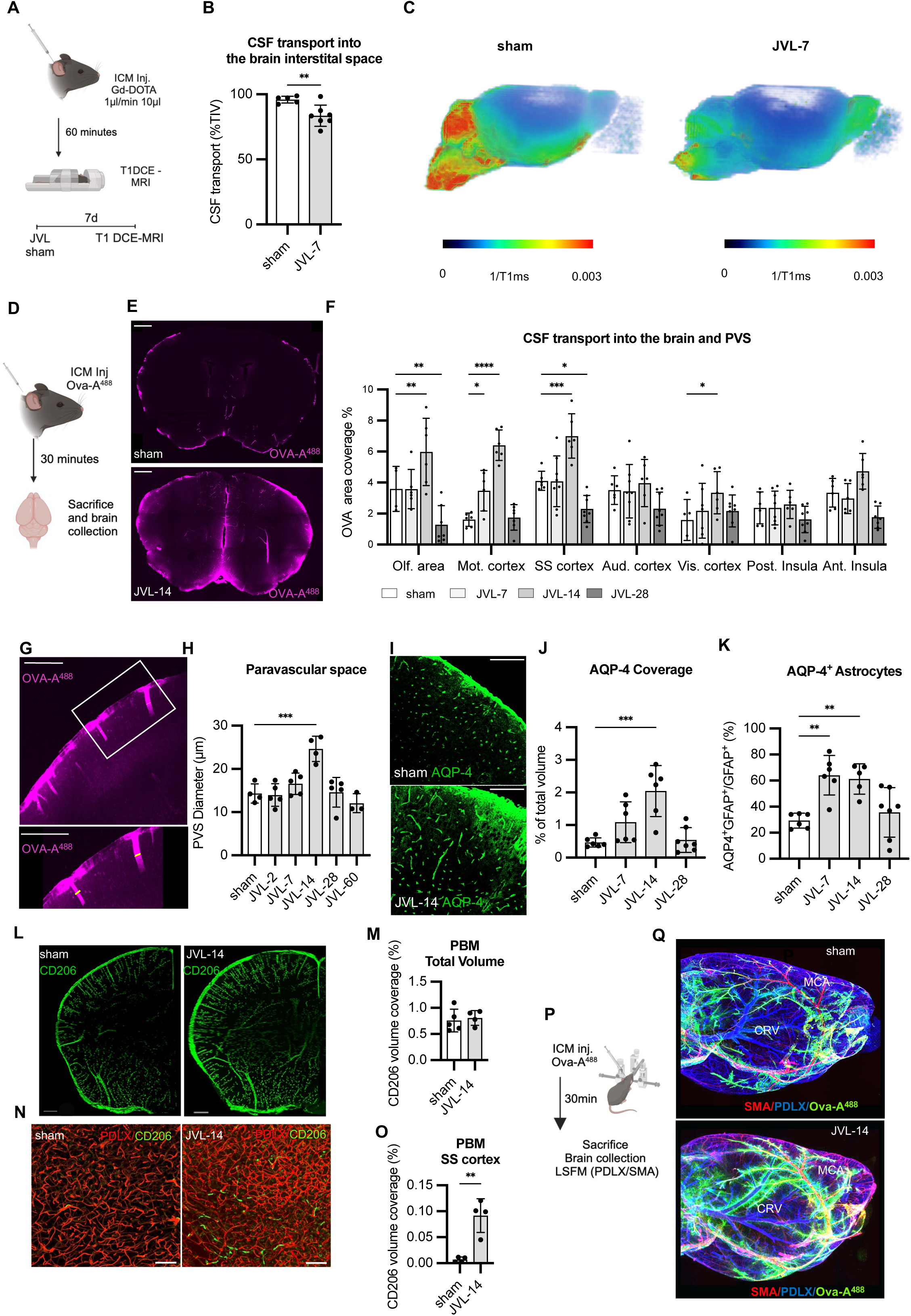
Brain clearance kinetics after jugular vein ligation. (A, C) Intracerebral drainage of CSF. Scheme of experimental design, using T1 dynamic contrast-enhanced (DCE) MRI after ICM. injection of Gd-DOTA in sham operated and JVL mice at 7-dps (A). MRI scans show that jugular ligation strongly reduced diffusion of contrast agent into the brain parenchyma (B-C), as quantified in (B). (D, F) Assessment of intracerebral CSF distribution. Experimental design: CSF in perivascular spaces (PVS) and brain tissue was evaluated at 30 minutes after ICM. injection of OVA-A^488^ in sham operated and JVL mice (D). OVA-A^488^-labeled area on brain slices at the level of the somatosensory cortex (SS) in mice at 14-dps (E). Quantification of OVA-A^488^ labeling showed that, in JVL mice, CSF entry was increased in both the PVS and parenchyma of all cortical regions draining into the SSS, including olfactory (Olf.), motor (Mot.), somatosensory (SS), and visual (vis.) cortex with non-significant trends in the auditory (aud.) cortex and insula (F). (G, H) PVS enlargement at 14-dps after jugular ligation. At 30 min after ICM. injection of OVA-A^488^, mice were sacrificed, and labeled PVS were measured on brain sections (G). Quantification shows enlargement of PVS diameter at 14-dps (H). (I, K) Kinetics of AQP-4 pattern after jugular ligation. Confocal imaging of the SS cortex of sham operated and JVL-14 mice after immunolabeling with anti-AQP-4 antibodies (I). Volumetric quantifications show increased AQP-4 coverage at 14-dps (J). Increased expression by astrocytes (AQP-4^+^GFAP^+^) was observed at 7- and 14-dps in the SS cortex. (L, O) Pattern of PBM at 14-dps after jugular ligation. (L, M) CD206^+^ PBMs imaging by LSFM in forebrain sections of sham operated and JVL-14 mice (L) revealed that the global PBM population was not changed by jugular ligation. (N, O) CD206^+^ PBMs of the SS cortex. Confocal imaging of CD206^+^ PBM in the SS cortex of sham operated and JVL-14 mice (N). Quantitative assessment of the increased PBM coverage along cortical vessels in JVL-14 mice (O). (P, Q) Paravenous concentration of phagocytic cells in JVL mice at 14-dps. Experimental design: ICM. injection of OVA-A^488^ (green) at 30 minutes prior to sacrifice, treatment of whole brains with iDISCO^+^ and immunolabeling of PDLX^+^ (blue)/SMA^+^ (red) arteries and PDLX^+^/SMA^-^ veins, and imaging with a LSFM (P). Lateral volumetric views of the forebrain of sham operated and JVL-14 mice (Q). Numerous OVA-A^488^-labeled cells are observed in the paravascular spaces of JVL-14 mice compared to controls. Note that phagocytic cells surround the middle cerebral artery (MCA, SMA^+^/PDLX^+^) as well as the caudal rhinal vein (CRV, SMA^-^/PDLX^+^) in JVL mice. Only few phagocytic cells are detected in the paravenous spaces of control mice. Two-way ANOVA with Dunnett’s multiple comparisons test was used in F. Ordinary one-way ANOVA with Dunnett’s multiple comparisons test was used in H, J, and K. Unpaired t-test was used in B. Mann Whitney U-test was used in M, and O. \**P* < 0.05, \*\**P* < 0.01, \*\*\**P* < 0.001, \*\*\*\**P* < 0.0001. Error bars indicate SD. Scale bars: 1000 µm (E), 500 µm (G, L), 200 µm (I), 100 µm (N). Elements in A, D and P were created using BioRender (https://biorender.com).

We reasoned that intracerebral CSF drainage may resume to normal once dural lymphatic vasculature was restored at 14-dps. After ICM injection of Ovalbumin-Alexa (OVA-A^488^) to visualize intracerebral CSF drainage (Fig. 4D), we found enlarged paravascular spaces (PVS) with increased permeability as attested by increased OVA-A^488^ coverage in the cortical regions in JVL mice at 14-dps (Fig. 4E-H). The diameter of paravascular spaces increased along with the astroglial expression of aquaporin 4 (AQP-4) (Fig. 4I, J), which contributes to fluid exchanges between paravascular spaces and the brain parenchyma. ^12^ Notably, the number of astrocytes expressing AQP-4 increased at an earlier timepoint, 7-dps, suggesting that the mechanisms underlying PVS remodeling were initiated prior to 14-dps (Fig. 4K).

Paravascular CSF drainage depends on parenchymal border macrophages (PBMs).^24^ Using volumetric imaging by light-sheet fluorescence microscopy (LSFM), we assessed the total volume of PBMs in sham operated and JVL-14 mice (Fig. 4L). While the total PBM volume did not differ significantly between these two groups (Fig. 4M), PBMs in JVL-14 mice appeared to penetrate deeper into brain tissue. To confirm this, we used confocal microscopy to quantify CD206^+^ PBMs along cerebral vessels in the somatosensory cortex of JVL mice. As expected, PBM numbers increased at 14-dps, aligning with enhanced intracerebral CSF drainage (Fig. 4N, O). We further examined paravascular phagocytic cells through LSFM and intracerebral injection of fluorescent OVA-A^488^ (Fig. 4P), along with immunolabeling for podocalyxin (PDLX) and smooth muscle actin (SMA) to distinguish arteries (PDLX^+^/SMA^+^) from veins (PDLX^+^/SMA^-^). In JVL-14 mice, we observed accumulation of OVA-A^488+^ labeled phagocytic cells within cortical paravenous spaces, whereas these spaces were sparsely labeled in sham operated mice (Fig. 4Q). This data shows that active remodeling of astroglial and PBMs along paravascular spaces accompanies the restoration of intracerebral CSF drainage post-JVL.

### Dural lymphatics regulate intracerebral pressure and CSF drainage

To determine whether dural lymphatic remodeling contributes to changes in brain fluid clearance following JVL, we examined the effects of enhanced or reduced dural lymphatic function on intracerebral pressure, perivascular CSF flow, and lymphatic drainage in sham operated and JVL mice. ICM injection was used to deliver either AAV-VEGF-C to promote dural lymphatic drainage, or AAV-VEGF-C trap (VEGFR-3_1-4_-Ig) to induce regression of dural lymphatics, as previously described.^7,25–27^ AAV-treated mice underwent sham or JVL surgery four weeks after ICM injection, followed by ICeP measurement prior to sacrifice (Fig. 5A). We predicted that impaired lymphatic function would exacerbate the effects of JVL and prolong the elevation of ICeP; therefore, we assessed ICeP in VEGF-C trap-treated mice at 7-dps. Conversely, we hypothesized that enhancing lymphatic function would facilitate CSF drainage after JVL, so we measured ICeP in AAV-VEGF-C-treated mice at 2-dps. Both sham operated and JVL mice with impaired dural lymphatics had increased ICeP (Fig. 5B, C). Notably, dural lymphatic ablation alone increased ICeP without additional elevation from venous ligation, indicating that JVL raises ICeP primarily through the regression of dorsal lymphatics. Prophylactic enhancement of dural lymphatic drainage using AAV-VEGF-C in JVL mice slightly reduced ICeP, though it remained significantly elevated compared to controls (Fig. 5D). In mice with impaired lymphatics, at 14-dps, intraparenchymal CSF circulation showed no recovery, as indicated by similar OVA-A diffusion and paravascular space diameter compared to sham operated mice (Fig. 5E-G). Together, these findings demonstrate that ICeP regulation and brain clearance depend on dural lymphatic drainage. We further confirmed that loss of dorsal lymphatics observed at 7-dps after JVL contributed to accelerated CSF drainage into cervical lymph nodes, as lymph node drainage in AAV-VEGF-C trap-treated mice was also accelerated compared to controls (Fig. 5E). In summary, dural lymphatic drainage directly regulates brain clearance and ICeP, with cerebral venous blood flow indirectly supporting these processes by maintaining dural lymphatic integrity.

**Figure 5.**
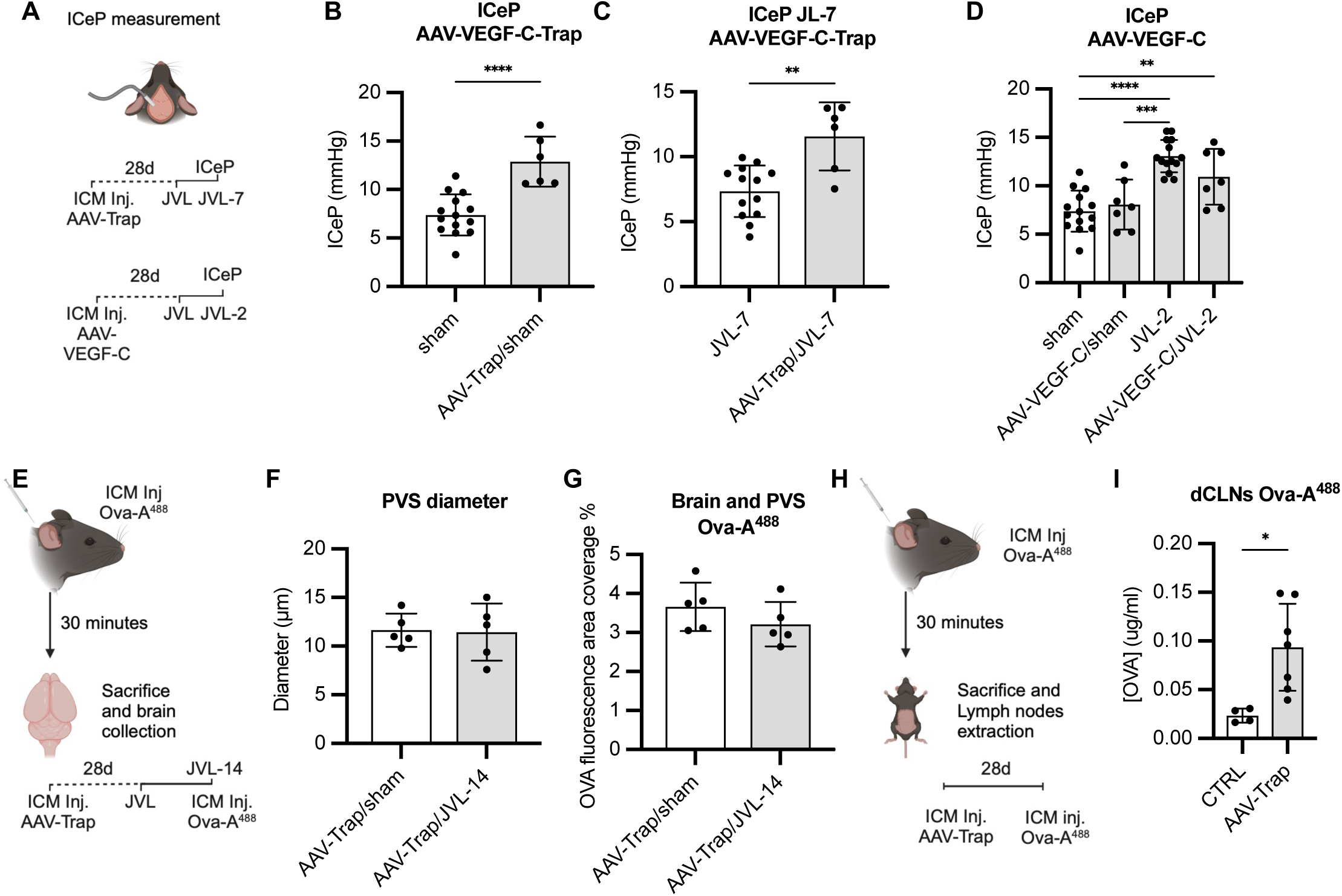
Dural lymphatic regulation of ICeP and brain clearance. (A, D) Effect of dural lymphatics manipulation on intracerebral pressure (ICeP). Intracerebral pressure (ICeP) was measured in sham operated and JVL mice previously treated to block or enhance dural lymphatic function, by ICM. injection of either AAV-VEGF-C trap or AAV-VEGF-C, respectively (A). Dural lymphatic loss raised ICeP in non-ligated mice (B) and impaired the restoration of ICeP at 7-dps in JVL mice (C). Enhanced lymphatic function does not change ICeP in control mice and does not significantly reduce ICeP at 2-dps in JVL mice (D). (E-G) Intracerebral CSF drainage in dural lymphatic deficient-JVL mice. Experimental design: sham operated and JVL mice pre-treated with AAV-VEGF-C trap received ICM injection of OVA-A^488^ at 14-dps (E). Quantification of intracerebral CSF flow in the paravascular spaces (F) and the parenchyma (G) of sham operated and JVL mice pre-treated with AAV-VEGF-trap at 14-dps. Sham operated mice without ligation are used as baseline controls. In contrast to JVL mice (Fig. 4H), no enlargement of PVS was detected in dural lymphatic-depleted mice, without or with jugular ligation, compared to controls (F) and no increase in CSF transfer into the cerebral parenchyma was observed in dural lymphatic-depleted mice (G). (H, I) dcLN drainage in JVL mice with loss of dural lymphatics. OVA-A^488^ content in dcLNs was quantified 30 min after ICM. injection of OVA-A^488^ (H). dcLN drainage of tracer increased in JVL mice with deficient dural lymphatics, compared to JVL controls (I). Unpaired t-test was used in B, C, F, G and I. Ordinary one-way ANOVA with Tukey’s multiple comparisons test was used in D. \**P* < 0.05, \*\**P* < 0.01, \*\*\**P* < 0.001, \*\*\*\**P* < 0.0001. Error bars indicate SD. Elements in A, E and H were created using BioRender (https://biorender.com).

### Lymphatic restoration in JVL-14 mice is driven by dural mesenchymal and endothelial cells interactions

To gain molecular insights into the mechanisms of lymphatic restoration observed in JVL-14 mice, we performed scRNA-seq analysis on dura mater isolated from JVL-14 and sham operated mice (Fig. 6A and SF. 4A). Clusters were annotated by comparing gene expression profiles to reference marker datasets (Fig. 6B and SF. 4B, C).^28,29^ Analysis focused on blood and lymphatic endothelial cells (hereinafter referred as ECs) and mesenchymal cells (SF. 4D, E). Gene set enrichment analysis (GSEA) indicated that the Fibroblasts-1 cluster displayed the highest number of signaling pathway modifications after JVL (SF. 4F), while the cluster of ECs showed enrichment in signaling pathways regulating the translation machinery, like eukaryotic translation initiation (n = 112; p = 0.01502) and elongation (n = 86; p = 0.01502), as well as in Slit/Robo signaling genes (n = 147; p = 0.02296) which participate in developmental angiogenesis and vascular injury repair ^30–32^ (Fig. 6C). The Fibroblasts-1 cluster of JVL-14 mice presented anti-inflammatory signatures, as indicated by the downregulation of interferon gamma response (n = 150; p = 0.01544), TNFα signaling via NF-κB (n = 150; p = 0.04476), signaling by interleukins (n = 283; p = 0.02105) and cytokine signaling in immune system (n = 441; p = 0.02185), while the TGFβ signaling pathway was upregulated (n = 46; p = 0.04476) (Fig. 6D). We next analyzed the differential ligand–receptor (L–R) interactions between clusters (Fig. 6E). The interactions between ligands produced by Fibroblasts-1 and receptors expressed by ECs was increased in JVL-14 mice, with reinforced VEGF-C-VEGFR-2/3 and Angiopoietin-2 (Ang-2) signaling interactions, suggestive of active dural lymphangiogenesis (Fig. 6F, green asterisks). Especially, *Cdkn1a* gene, also known as *p21* and a negative regulator of cell cycle acting downstream of growth factor receptors ^33,34^, including VEGFR-2 was highly downregulated in dural ECs of JVL-14 mice (logFC = -1.76460; p = 4.27999 x 10^-6^) (Fig. 6G). The possible contribution of VEGF-C in dural lymphatic remodeling after JVL was moreover supported by the increased level of VEGF-C measured in the plasma from 7-dps (Fig. 6H). These findings correlate with the regrowth of dural lymphatics observed in JVL-14 mice and suggest that subsets of fibroblasts of the dura mater may drive lymphangionic repair.

**Figure 6.**
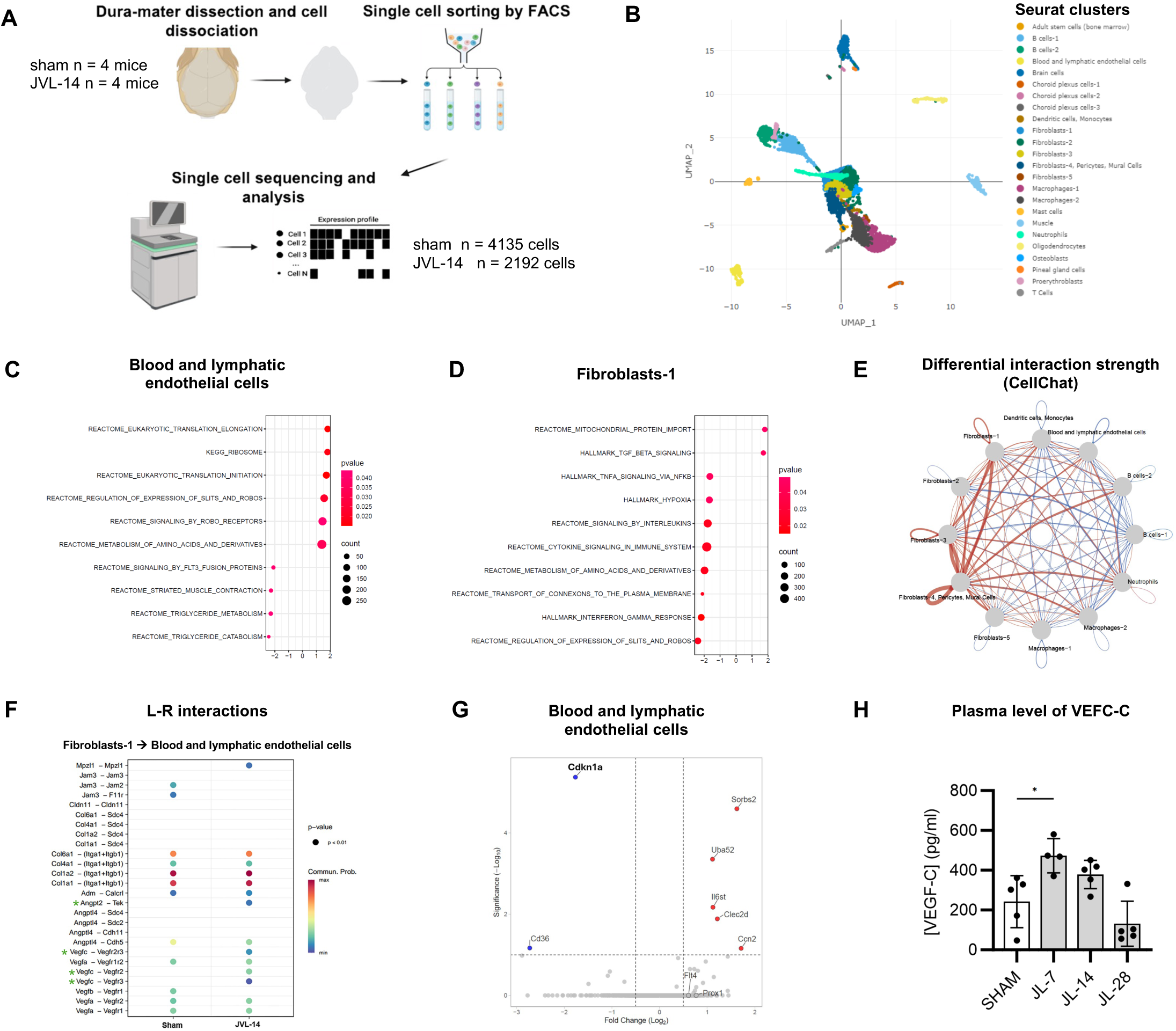
scRNA-seq analysis of dura mater in JVL-14 and sham operated mice. (A) Scheme of experimental design (n = 4 mice / group). (B) UMAP showing identified clusters within merged data of sham operated and JVL-14 dura. (C, D) Dot plots from Gene Set Enrichment Analysis (GSEA) highlighting selected significantly regulated pathways in mesenchymal and vascular endothelial cells of JVL-14 mice compared to sham operated mice. Slit/Robo signaling pathways and translation signatures were enriched in Blood and lymphatic cells (C) and anti-inflammatory signatures were enriched in Fibroblasts-1 clusters (D). (E) Differential interaction strength of CellChat ligand-receptor (L-R) interactions between selected cell types. Red and blue lines indicate upregulated and downregulated interactions, respectively, in JVL-14 mice. (F) Fibroblasts-1 cluster ligands exhibit increased communication probability with Blood and lymphatic endothelial cells cluster receptors mediating VEGF-C lymphangiogenic signaling (green asterisks) in JVL-14 mice compared to sham operated mice. (G) Volcano plot of up- and down-regulated genes in the Blood and lymphatic endothelial cell cluster. (H) Monitoring of the plasma level of VEGF-C over time after jugular ligation, showing overexpression of VEGF-C in JVL-7 mice. Selected regulated pathways with an adjusted p-value ≤ 0.05 are plotted (D, E). Endothelial and mesenchymal cell clusters and immune cells clusters containing at least 50 cells in both conditions were included in the analysis (F). L-R interactions with significant differences in communication probability between conditions are displayed (G). Ordinary one-way ANOVA with Dunnett’s multiple comparisons test was used in H. \**P* < 0.05. Error bars indicate SD. Scheme in A was created using BioRender (https://biorender.com).

## Discussion

The present study of cerebral blood flow and CSF drainage in patients with IIH and JVL mice shows that the dural veno-lymphatic system regulates brain clearance. In both models, we correlated the impairment of cerebral venous outflow with the regression of calvarial lymphatics in the dura mater and the accumulation of extravascular fluids in the brain. Brain oedema contributed to intracranial hypertension, although without ventricular enlargement. We concluded that cerebral blood venous outflow regulates the maintenance of calvarial lymphatics and brain fluid drainage.

In patients with IIH, brain oedema and intracranial hypertension are consistently associated with the presence of venous stenoses and regressed perisinusal fluid drainage in the transverse sinuses. Parasagittal fluids accumulate in the dura along the superior sagittal sinus as well as along the bridging veins in the subarachnoid space. Evidence for a structural and functional continuity of cerebral paravenous spaces with the dural perisinus comes from DCE-MRI data obtained with the 3D T1 SPACE DANTE sequence in IIH patients. Contrast enhancement was simultaneously observed along cortical bridging veins and around the superior sagittal sinus (Fig. 1G). In the pathological condition of IIH, the paravenous CSF tracer is therefore in relation with the dural perisinusal space. ^35^ On one hand, this observation suggests that subarachnoid, and by extension cerebral paravenous fluids, accumulate in the dural perisinus, aligning with the model proposed by Smyth et al., which describes specialized channels along bridging veins, referred to as arachnoid cuff exit (ACE) points, that facilitate fluid and immune exchanges between the subarachnoid space and the dural perisinusal space.^15^ On the other hand, the detection of contrast agent in subarachnoid paravenous spaces may indicate a reflux of dural fluids accumulated in excess in the dura mater of IIH patients, as a result of the blockade of lymphatic drainage in the transverse and sigmoid sinuses. Future experiments by dynamic imaging of paravenous and lymphatic drainage will allow to state about this alternative. A third explanation for the accumulation of contrast agent in the parasinus is a possible leakage of venous fluid into the perisinus due to sinus hypertension. While we did not observe leakage of injected tracers at the venous sinuses in mice even at the peak of intracerebral hypertension post-JVL, we cannot exclude such leakage may occur in IIH patients (SF. 2D). However, it is equally possible that the presence of contrast agent in the perisinus reflects the defective drainage of paravenous fluids via downstream lymphatics.

The drainage of perisinusal fluids into the dural lymphatics of the calvaria therefore remains the most plausible explanation of the phenotype observed in IIH patients. Venous stenosis was associated with depletion of lymphatics in the transverse sinus (TS), while perisinusal space was enlarged along the superior sagittal sinus (SSS). The flow of perisinusal fluid from the SSS to the TS is likely blocked by the interruption of TS lymphatic-like tubules which are located downstream of the SSS enroute to the cervical lymph nodes. This novel observation supports the role of dural lymphatics as an exit route of brain drainage fluids in humans. The presence of brain oedema in IIH patients agrees with the previous report that glymphatic outflow is blocked in IIH patients.^36^ Radiological dilation of cranial nerve sheaths ^37^ combined with regression of calvarial lymphatics suggest that CSF clearance may proceed mainly through the perineural routes in these patients. Further studies will be required to confirm fluid transfer from the perisinusal spaces to dural lymphatics. Non-human primates, with calvarial lymphatics positioned similarly to those in humans, provide an ideal model since they are amenable to both dural sinuses interventional surgery and DCE-MRI analysis of dural lymphatics.^8,14^ The cause of dural lymphatic regression can only be speculated but may be directly related to the increase of intravenous pressure occurring in the dural sinuses of IIH patients. ^38^ Within the rigid envelope of the dura mater, the physical proximity of lymphatics with venous sinuses under hypertension may cause compression of lymphatics and impair lymphatic drainage along the SSS, leading to regression of downstream lymphatics. The stenting of TS enables normalization of venous sinus flow and intracranial pressure in IIH patients. ^2^ Further MRI studies will be required to assess whether venous sinus stenting allows restoration of calvarial lymphatic flow as predicted.

Blockade of cerebral outflow in mice was induced by surgical ligation of jugular veins. Like IIH patients, JVL mice showed regression of calvaria lymphatics, abnormal CSF drainage, and intracerebral hypertension. However, in contrast to IIH patients, the above-mentioned phenotype was transient. A complex remodeling of the blood and lymphatic vasculature coordinated with plastic changes in cerebral paravascular spaces in the JVL mice, which thus provide a potential model for mechanistic studies of brain vascular remodeling. Occlusion of venous blood flow caused regression of dural lymphatics in the calvaria, as well as a profound remodeling of the arterial vasculature. In return, restoration of dural lymphatics at two weeks after JVL synchronized with a normalization of cortical blood perfusion. The correlation observed between vascular remodeling and fluctuations of brain fluid homeostasis underscores the critical role of cerebral blood flow and dural lymphatic drainage for cortical fluid homeostasis. During the first week after the surgery, the acute peak of intracerebral hypertension at 2-dps was followed by the regression of calvarial lymphatics and the blockade of CSF entry into the brain. The closure of the glymphatic influx was thus synchronized with the loss of lymphatic efflux in response to the blockade of cerebral venous outflow. During the second week after jugular ligation, lymphatic recovery in the dura mater correlated with plastic remodeling of cerebral blood vessels and perivascular spaces, resulting in enhanced perivascular CSF drainage. Re-opening of lymphatic efflux synchronized with reactivation of glymphatic efflux and re-perfusion of CSF along perivascular spaces. These observations indicate an interplay between dural lymphatic activity and intracerebral CSF drainage. Evidence for a causal role of dural lymphatics in brain clearance is supported by three findings resulting from the ablation of dural lymphatics by ICM injection of VEGF-C trap. Specifically, ablation (1) caused cerebral hypertension in control mice; (2) resulted in maintained cerebral hypertension after JVL while it was restored at 7-dps in control JVL mice; and (3) blocked the re-perfusion of CSF along perivascular spaces observed in JVL-14 mice. Brain fluid homeostasis is, therefore, dependent upon dural lymphatic drainage. The regulatory mechanisms of this interplay between the glymphatic and lymphatic systems may involve, at least in part, the population of parenchymal border macrophages which increased during the second week after ligation and has been already shown to regulate glymphatic drainage. ^24^ Transcriptomic analysis indicates that the dura mater may provide complementary regulation through interactions between mesenchymal and endothelial cells. The resulting modulation of VEGF-C/VEGFR-3 and Ang-2 signaling pathways, which are key lymphangiogenic regulators^25,39^, may explain the increased expression of plasma VEGF-C at 7-dps and the regrowth of calvaria dural lymphatics in JVL-14 mice. TGFβ is known to upregulate VEGF-C expression. ^40,41^ TGFβ signaling genes are upregulated in the Fibroblasts 1 cluster of JVL-14 mice, which may explain the increased VEGF-C expression by these cells. TGFβ expression is moreover regulated by mechanosensing^42,43^, suggesting that dural lymphangiogenic repair may result from sheer stress conditions developed in response to JVL. Taken together, our results demonstrate that the veno-lymphatic complex is a co-dependent vascular system essential for brain clearance. Cerebral venous blood flow indirectly influences intraparenchymal CSF circulation by maintaining calvarial lymphatics, which appears to be the primary route for cortical CSF drainage.

## Material and Methods

### Study approval

Human subjects: The prospective Lymph-IMAGIIH study was approved by our institutional review board (IDRCB: 2022-A01422-4) and registered on ClinicalTrials.gov (ID: NCT05762367), with INSERM (Institut National de la Santé et de la Recherche Médicale) serving as the study sponsor. Mice: All in vivo procedures complied with guidelines and institutional policies of INSERM and the Animal Care and Use Committee of the Paris Brain Institute (APAFIS#34974-2022012017455126).

### Human subjects

The Lymph-IMAGIIH study included two groups of female participants aged 18–40 years: 18 patients with idiopathic intracranial hypertension (IIH) and 20 age- and sex-matched healthy controls. Among the IIH group, 16 had complete MRI scans; one participant was unable to undergo MRI due to claustrophobia, and intravenous (iv.) gadobutrol injection failed in another one. No significant age difference was observed between groups (healthy controls: 28.0 (4.96) years; IIH patients: 30.2 (4.88) years). IIH patients had a higher body mass index than controls (31.6 (9.49) vs. 23.1 (3.98); p < 0.001). The mean CSF opening pressure in IIH patients was 32.4 (4.24) cm H₂O. Fourteen patients were on acetazolamide at the time of MRI, with a mean dosage of 1107 (691.4) mg/day.

### Human MRI protocol

All radiological data were pseudonymized, with postprocessing conducted by an experienced neuroradiologist and a neurologist. MRI acquisitions were performed on a 3.0 Tesla Magnetom Skyra (Siemens Healthineers) in a single session under 45 minutes. The complete MRI protocol was achieved in 16 subjects with IIH and in all 20 healthy volunteers. Details of the sequences are as follows:

Pre-gadobutrol injection: T1w MPR1 iso, 3D Sagittal acquisition, 224 mm^2^ FOV, 256 contiguous 1.00 mm slices, TR/TE 2400.0/2.02 ms, 5.31 min acquisition time; 3D TOF, 3D transversal, 230 mm^2^ FOV, contiguous 0.40 mm slices, TR/TE 21.0/4.44 ms, 4.04 min acquisition time; T2 SPC FLAIR ISO1 real, 3D Sagittal, 196 mm^2^ FOV, contiguous 1.00 mm slices, TR/TE 15230/549.0 ms, 6.36 min acquisition time.

Post-gadobutrol injection (gadovist^®^, 0.1 mmol/kg; molecular weight: 558.64 g/mol)):

Contrast-enhanced MR venography, elliptical-centric, 3D Coronal, 250 mm^2^ FOV, contiguous 0.70 mm slices, TR/TE 3.57/1.36 ms, 1:20 min acquisition time; 3D SPACE FLAIR, 3D Coronal, 256 mm^2^ FOV, contiguous 1.00 mm slices, TR/TE 5000/375 ms, 5.15 min acquisition time; 3D SPACE T1 DANTE, 3D Sagittal, 307 mm^2^ FOV, contiguous 0.80 mm slices, TR/TE 700/21.0 ms, 6.33 min acquisition time; T2 SPC FLAIR ISO1 real, 3D Sagittal, 196 mm^2^ FOV, contiguous 1.00 mm slices, TR/TE 15230/549.0 ms, 6.36 min acquisition time.

### Post-processing of human MRI scans

MRI scans were processed using the 3D Slicer platform, employing semi-automated thresholding for signal intensity and segmentation of native sequences as previously described by Jacob et al.^44^ The venous system was segmented from the Venous Elliptical sequence, the lymphatic system from the 3D T1-SPACE DANTE sequence, and brain, ventricular and choroid plexuses structures from the FLAIR sequence. Total intracranial volume was derived from the 3D T1 MPRAGE sequence. Additionally, the choroid plexus was segmented based on the FLAIR sequence using automatic segmentation, as previously described by Yazdan-Panah et al.^45^

### Statistical analyses of human data

Independent measurements were performed by a neuroradiologist and a neurologist, both blinded to pathology, to assess reproducibility. Inter-rater agreement was evaluated using the Intraclass Correlation Coefficient (ICC), with a two-way random model and absolute agreement. ICC values were as follows: brain volume (0.79), total intracranial volume (0.83), ventricles volume (0.69), perisinusal volume (0.92), and venous volume (0.75). All variables met normality assumptions, as confirmed by Shapiro-Wilk and Levene’s tests. Data were expressed as mean (SD), with error bars representing SD. Group differences were therefore analyzed using a t-test, with statistical significance set at p < 0.05.

### Mice

C57/BL6 mice, aged 2–4 months, were obtained from Jackson Laboratories and housed under specific pathogen-free conditions. All experiments were performed in female mice, as the conditions of dural venous stenosis and idiopathic intracranial hypertension predominantly affect women in humans, except for two-photon imaging, which was conducted in adult male B6.129P2(Cg)-Cx3cr1GFP mice. The transgenic Prox1-Cre-tdTomato reporter mice used in this study were kindly provided by Dr. Steven T. Proulx at the University of Bern, Switzerland. Two-photon imaging was performed at the Institut Pasteur, Paris, France.

## Surgery procedures

### Jugular vein ligation

To perform cerebral venous outflow obstruction, adult C57/BL6 female mice underwent bilateral ligation of the internal and external jugular veins. Prior to surgery, mice received an intraperitoneal injection of buprenorphine (0.1 mg/kg) and carprofen (20 mg/kg), along with 200 μL of 0.9% sodium chloride for hydration. Anesthesia was induced with isoflurane (3–4% at 400–500 mL/min airflow) and maintained at 1.5% (140–250 mL/min, adjusted to respiratory rate). Mice were positioned supine on a heated pad to maintain body temperature at 37°C, monitored via rectal probe. The surgical area was disinfected, and aseptic conditions were upheld. A longitudinal incision was made in the ventro-cervical region to expose the jugular veins, which were then ligated with 6.0 silk sutures. Skin incisions were closed using non-absorbable propylene sutures. Sham operated mice underwent the same exposure procedure without vein ligation.

### Intracerebral pressure (ICeP) measurement

Following anesthesia induction (3-4% Isoflurane, maintenance at 1.5% with adjusted airflow), a 1 cm longitudinal skin incision was made to expose the skull. A 0.5 mm hole was drilled above the parietal lobe, and a pressure sensor catheter (SPR100, Millar) was inserted 1 mm into the cortex using a stereotaxic frame. ICeP was monitored for 6 minutes, and mean values were recorded over the last 3 minutes after stabilization.

### Intra-cisterna magna (ICM.) injection

Prior to surgery, mice received intraperitoneal injections of buprecare (0.1 mg/kg) and carprofen (20 mg/kg) for analgesia, and 200 μL of 0.9% sodium chloride for hydration. Following anesthesia induction (3-4% Isoflurane, maintenance at 1.5% with adjusted airflow), mice were positioned on a stereotaxic frame with the head tilted at a 120° angle. A 1 cm neck incision exposed the cisterna magna, which was punctured with a 27G needle. We used a Hamilton syringe (2 µL, 30 gauge) connected to a Picospritzer III precision injection system inserted at a 90° angle to deliver ICM. injections of ovalbumin conjugated with either AlexaFluor™ 488 (OVA-A^488^) or AlexaFluor™ 647 (OVA-A^647^) (10 ml, 2 mg/mL, 2 µL/min). AAV_9_ vectors expressing VEGF-C or VEGFR-3_1-4_-Ig (VEGF-C trap) were delivered at titers between 3.53 × 10¹¹ and 6.18 × 10¹^2^ pv/mL, in 2 µL per mouse at 0.5 µL/min. The neck incision was thereafter sealed with surgical glue.

### ICM. injection of fluorescent tracers with a delivery catheter

Prior to surgery, mice received intraperitoneal injections of buprecare (0.1 mg/kg) and carprofen (20 mg/kg) for analgesia, and 200 μL of 0.9% sodium chloride for hydration. Following anesthesia induction (3-4% Isoflurane, maintenance at 1.5% with adjusted airflow), the surgical site was shaved, disinfected, and the mouse was placed ventrally on a heated pad. 2 cm neck incision exposed the cisterna magna, where a catheter was implanted at a 30° angle. The incision was sealed with surgical glue. Two injections of different fluorescent tracers (OVA-A^647^ and OVA-A^488^, 4 mg/mL) were administered through the catheter: the first immediately after catheter placement (120 minutes prior to euthanasia) and the second 30 minutes prior to euthanasia. Each injection delivered 2 μL at 1 μL/min. Mice were awakened between injections.

### In vivo two-photon imaging

Mice were anesthetized (100 mg kg^-1^ ketamine/10 mg kg^-1^ xylazine), the dorsal skull was exposed, and a 3-mm square cranial window was made over the right somatosensory/motor cortex.^46^ Dental cement was used to seal both the cover-glass and to secure a titanium custom head-bar (0.5 g) for attaching to a custom stage. After >4 weeks, under isoflurane anesthesia, the mice had a retro-orbital injection of 70k MW dextran-tetramethylrhodamine (50 μl, 5 mg ml^-1^, D1818, Invitrogen, Waltham, MA, USA) to label blood vessels. For *in vivo* 2-photon imaging, the mice were lightly anesthetized with isoflurane (0.8%, humidified oxygen), the head-bar was secured to the custom stage and body temperature was maintained with a rectal thermometer feedback heating pad. The mouse stage was zeroed to the microscope with a fluorescent fiducial reference for using coordinates to find the same fields of view for each imaging session. Imaging was performed with an upright microscope (Investigator microscope, Bruker, Madison, WI, USA) with a 16x objective (CF175 LWD 16X W, 0.8 NA, Nikon Instruments, Tokyo, Japan) using a Ti:Sapphire laser (Mai Tai DeepSee, Sectra Physics, Milpitas, CA, USA) tuned to 840 nm. For each imaging session, a z-stack (0.483 μm/pixel x-y, 3 μm z-step) was acquired and line scans (∼4 kHz) were performed along the center of the long-axis of vessels within the reference brain volume for blood flow measurements. ^47^

### Two-photon blood vessel analysis

Line scan reference images were registered into the imaging session z-stacks for identifying line scanned vessels and the scan direction. Blood flow direction for each scanned vessel was annotated with arrows (FIJI-ImageJ, NIH) and vessel types were segregated between artery and vein categories based on 1) surface to deep flow for arterial structures and 2) converging flow from deep to surface for veinous structures. Using the directionality analysis plugin^48^ (“Fourier components” method, FIJI-ImageJ) the trajectory angle of line scan images (x-distance vs. y-time) was detected. The blood flow was calculated:

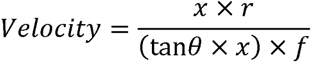

With the line scan length (*x*), scan resolution (*r*), trajectory angle (*θ*), and line scan frequency (*f*). Vessel diameter was measured at the same anatomical location within the z-stack volume across all timepoints and the data were plotted for statistical analyses.

## MRI procedures and postprocessing of images

### MRI after intravenous (IV.) injection of gadobutrol

MRI scans were performed on a preclinical 11.7 T MRI scanner (Biospec Bruker, BioSpin, Germany) equipped with a 1H surface cryoprobe for mouse brain imaging. A metal-free, MRI-compatible catheter (MTV-33, 28 cm PU, 1F, 29G) was inserted into the tail vein and connected to a syringe pump (model 70-2130, Harvard Apparatus) preloaded with gadobutrol (gadovist, 1.0 mmol/mL). The system was purged and flushed with contrast agent prior to use. Mice were anesthetized with isoflurane (4% induction, 1–2% maintenance, 500 mL/min oxygen flow) and monitored for respiratory rate, while body temperature was maintained at 37 ± 1°C. Heads were positioned supine and secured using a bite bar and ear pins. Gadobutrol (400 μL at 50 μL/s) was injected automatically 20 seconds before the T1-weighted perfusion sequence. MRI Protocols Included 2D Time-of-Flight (TOF) (TE = 3.13 ms, TR = 24.4 ms, FA = 80°, BW = 125 kHz, 80 slices, slice thickness = 300 µm, FOV = 15.36 × 15.36 mm^2^, in-plane resolution = 60 × 60 µm^2^, acquisition time = 16 min 40 s), T1-weighted Perfusion (DCE-FLASH): TE = 1.2 ms, TR = 38 ms, FA = 15°, BW = 125 kHz, 10 slices, slice thickness = 800 µm, FOV = 12.8 × 14.8 mm^2^, spatial resolution = 200 × 200 µm^2^, acquisition time = 11 min 43 s. 3D Multi Gradient Echo (MGE): TE1 = 1.75 ms, ΔTE = 1.8 ms, TR = 40 ms, FA = 15°, BW = 200 kHz, FOV = 19.2 × 14.4 × 9.6 mm³, isotropic resolution = 100 µm, acquisition time = 9 min 13 s.

### Post-processing of DCE-MRI scans after IV. injection of gadobutrol

Anatomical region segmentations were performed using 3D-Slicer (https://www.slicer.org). The venous system was segmented from the 2DTOF sequence, while the brain and ventricles were segmented using the T1-MGE sequence. DICOM images from DCE-MRI acquisition were compiled into a 4D stack using Fiji ImageJ and converted to a NIfTI-1 file. To quantify perfusion, signal intensities of the tracer in the tissue and a feeding vessel were analyzed, facilitating the creation of time-dependent concentration curves and the application of a pharmacokinetic model to estimate quantifiable parameters. ImageJ was used to create masks for Regions of Interest (ROIs) and for Arterial Input Function (AIF) at the superior sagittal sinus (SSS). Quantitative analyses utilized a custom Python pipeline based on Python 3.11. The NiBabel library was used to read NIfTI-1 files, apply ROI and AIF masks, and extract signal intensities (S(t)) for each voxel at every time point. Pharmacokinetic modelling for parameter estimation was based on Mittermeier et al. (2019).^49^ For ROIs within the brain tissue, the signal intensity was converted to a time-dependent relaxation rate *R(t)* using the equation, where *S_0_* represents the initial signal intensity, α is the flip angle, and *TR* is the repetition time:

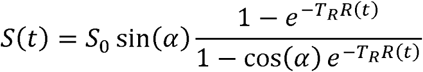

The tissue concentration *c_t_(t)* was calculated using specific relaxation rates of tissue *R1* and the gadolinium-based contrast agent *r1*:

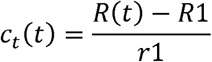

For the AIF, the same equations were used to estimate the tracer concentration in the selected vessel *c_b_(t)*. The hematocrit *hct* value of blood is used to convert blood tracer concentration *c_b_(t)* into concentration of tracer in the plasma *c_p_(t): c_p_*(*t*) = *c_b_*(*t*) · (1 - *hct*).

The tissue concentration curve *c_t_(t)* for each ROI and the AIF *c_p_(t)* were fitted into Toft’s and Kermode models using the tofts_integral and fit_tofts_model functions of the open-source Python module *DCEMRI*. Estimated K-trans values were extracted from the fitted models for each ROI.

### MRI with ICM. injection of Gd-DOTA for glymphatic transport imaging

MRI scans were performed on a Bruckner 9.4 T/16 MRI scanner, and T1 mapping was used to evaluate glymphatic transport in JVL and sham operated mice, after ICM. injection of gadoteric acid (Gd-DOTA (Guerbet LLC), 10 µl, 1 µl/min, molecular weight: 558 Da), as described in Boisserand et al.,2024.^20^ MRI experiments were conducted at the Magnetic Resonance Research Center at Yale University.

### Post-processing of Gd-DOTA MRI scans

Post-processing was performed as described in Xue et al., 2020.^22^ The T1 maps were filtered to exclude values greater than 5,000 ms. The MRIs, composed of summed low flip angle (2° and 5°) SPGR images, served as anatomical templates for outlining the brain and lymph nodes. Glymphatic transport volume was defined as brain tissue voxels with T1 values in the range of 1–1,700 ms, indicating tissue shortened by Gd-DOTA uptake. For each mouse, glymphatic transport volume was extracted using PMOD software (PMOD, version 4.0). Similarly, outflow to the nasal conchae was quantified by outlining these structures and extracting voxels with T1 values from 1 to 1,700 ms.

## Collection and analysis of biological material

### Plasma collection and Bioplex analysis

Blood was collected from mice via a cheek prick (0.2–0.3 mL) and treated with an intraperitoneal saline injection (0.9% NaCl; 10 mL/kg) for rehydration. Blood samples in EDTA tubes were centrifuged at 12,000 rpm for 5 min at 4°C, and plasma was stored at -20°C. Bioplex assays were performed using the MILLIPLEX® Mouse Angiogenesis/Growth Factor Magnetic Bead Panel - Cancer Multiplex Assay (MAGPMAG-24K-03, Merck KGaA, Darmstadt, Germany). at the ICM’s ICV3 platform, following the manufacturer’s protocol. Data analysis was conducted in Belysa® Immunoassay Curve Fitting Software (version 1.2.2, MilliporeSigma, Burlington, MA, USA, 2024).

### Tissue collection and processing

Following euthanasia mice were perfused with PBS and 4% paraformaldehyde (PFA). Brains, lymph nodes, skullcaps, and skull bases were harvested, fixed in PFA for 24 h, and stored in PBS. Portions of the brains and lymph nodes were cryoprotected in 30% sucrose, embedded in Tissue-Plus® O.C.T., and cryosectioned at 16 μm-thickness with a Leica cryostat, then stored at -20°C. The remaining tissues were kept in PBS and sectioned at 100 or 200 μm-thickness with a Leica VT1200 vibratome.

### Immunofluorescence labeling of whole-mount meninges

Decalcified skullcaps and skull bases (Surgipath Leica, 2 h) were blocked overnight in 0.3% Tween-20 and 5% donkey serum in PBS. Samples were incubated with primary antibodies (anti-LYVE1, anti-PDLX, anti-VwF) at 1/200 for skullcaps and 1/100 for skull bases in blocking solution for 3 days, followed by three 1-h PBS washes. Secondary antibodies (1/500 in 0.3% Tween-20 PBS) were applied for 48 h, followed by three additional 1-h PBS washes. Imaging was performed on a Leica Confocal SP8 X microscope with a 4x objective (5 μm Z-stack), and images were analyzed in ImageJ.

### Brain tissue preparation and analysis of OVA-A tracer distribution

Brains were extracted post-perfusion, fixed in 4% paraformaldehyde (PFA), and coronally sectioned at 100 μm using a Leica VT1000 S vibratome. Sections were mounted with Fluoromount™ Aqueous Mounting Medium. Glymphatic tracer distribution was analyzed on a Nikon AX R confocal microscope with NIS-Elements software. Regions of interest (ROIs) were delineated on DAPI-stained sections using Fiji ImageJ while utilizing the Allen Brain Atlas as a reference. For quantification of tracer in paravascular spaces and brain parenchyma, the Ovalbumin signal was binarized with a uniform threshold, and the percentage area of tracer coverage within each ROI was measured over time. Paravascular space diameter in the somatosensory cortex was determined by averaging width measurements from 5 to10 vessels on each ROI.

### Extraction and quantification of lymph node ovalbumin

Lymph nodes (deep cervical, superficial, lombo-aortic, sacral) were collected post-euthanasia and perfusion, placed in 2 mL tubes with 200 µL formamide, centrifuged at 10,000 g for 5 min, and incubated at 65°C for 48 h. An ovalbumin standard curve was prepared in formamide (20 to 0.0156 µg/mL). Samples (100 µL/well) were analyzed using a Spectramax® i3X plate reader (Molecular Devices).

### Immunofluorescence labeling on cryosections and free-floating brain sections

Cryostat brain sections were defrosted for 1 h, rehydrated in PBS for 15 min, and permeabilized in 0.5% Triton X-100 in PBS for 1 h, followed by three 5-min PBS washes. To prevent nonspecific binding, sections were blocked for 1.5 h at room temperature in 0.1% Tween-20 and 5% normal donkey serum in PBS. Free-floating sections were blocked overnight in 0.5% Triton X-100 and 5% normal donkey serum in PBS. Both cryostat and free-floating sections were incubated with primary antibodies overnight at 4°C. For cryostat sections, antibodies used were anti-Iba1 (1/150), anti-GFAP (1/100), and anti-AQP4 (1/100), and for free-floating sections, anti-CD206 (1/200), anti-PDLX (1/200), and anti-SMA (1/200). The next day, sections were washed in PBS and incubated with secondary antibodies (Alexa Fluor® 647, 555, and 488, diluted 1/500 for cryostat sections and 1/300 for free-floating sections in 0.1% or 0.5% Tween-20 PBS, respectively) for 2 h at room temperature for cryostat sections and overnight at 4°C for free-floating sections. Following three PBS washes, sections were stained with DAPI (10 µg/mL) for 2 min, then mounted with F4680 (Sigma-Aldrich) for cryostat sections or Mowiol for free-floating sections and stored at 4°C in the dark. Imaging was performed on a Nikon AX R confocal microscope with 3D segmentation using Nis Elements software.

### Tissue preparation and immunofluorescence labeling for iDISCO^+^

In iDISCO^+^ procedures with SMA (smooth muscle actin) - PDLX (podocalyxin) - CD206 labeling, and SMA - PDLX labeling after ICM.injection of OVA-A, samples were dehydrated through a methanol series (20%, 40%, 60%, 80%, and two 100% baths) at room temperature, followed by a dichloromethane (DCM)/methanol (2:1) bath for at least 4 hours until the samples sank. After bleaching overnight at 4°C in methanol containing 5% H₂O₂, samples were rehydrated through a reverse methanol series (80%, 60%, 40%, 20%) to PBS and washed twice in PTx.2 (PBS with 0.2% Triton X-100). Permeabilization was conducted in PTx.2 with DMSO and glycine for 2 days at 37°C, followed by blocking in a gelatin-based solution for 2 days. For the SMA-CD206 labeling, samples were incubated with primary antibodies, previously conjugated with Alexa Fluor™ Antibody Labeling Kits, for 10 days at 37°C with agitation, using rabbit anti-SMA (1:1000), goat anti-PDLX (1:1000), and goat anti-CD206 (1:1000). For the ICM Ovalbumin injection samples, primary antibodies, including rabbit anti-SMA (1:1000), goat anti-PDLX (1:1000), and goat anti-CD31 (1:500), were applied for 10 days at 37°C with agitation, followed by secondary antibodies (donkey anti-goat 555, 1:1000, and donkey anti-rabbit 790, 1:1000) for an additional 10 days. After PTwH washes (PBS with 0.2% Tween-20 and 10 µg/mL heparin), final dehydration was performed in a methanol series, followed by DCM/methanol (2:1) and two DCM baths, and then clearing in dibenzyl ether (DBE) until imaging.

### Light sheet fluorescence microscopy (LSFM)

Brains were imaged in the sagittal orientation using an LSFM (Ultramicroscope Blaze™, Miltenyi Biotec) with a 4× objective. Imaging was controlled by Imspector Microscope software (Version 7), with the chamber filled with DBE as a clearing agent. A one-sided, three-sheet illumination configuration was used with 2-step dynamic focusing. LED lasers at 561 nm, 640 nm, and 785 nm generated the light sheet, with a numerical aperture of 0.098. Emission filters included 595/40 for Alexa Fluor 594, 680/30 for Alexa Fluor 647, and 805LP for Alexa Fluor 785. Stacks were acquired with 3.5 μm z-steps, a 40-ms exposure time per step for the 561 nm and 640 nm lasers, and a 100-ms exposure time per step for the 785 nm laser. Mosaic acquisitions were performed with a 5% overlap on the full frame.

### Electron microscopy

Dura mater samples including venous sinuses and parasinus regions were fixed in a mixture of 2% paraformaldehyde, 2% glutaraldehyde, in 0.1 M sodium (Na) cacodylate buffer pH 7.4% overnight at 4°C, washed in 0.1 M cacodylate buffer and postfixed for 1h in 1% osmium tetraoxide in 0.1M Na cacodylate buffer. After washing in Na cacodylate buffer and water, samples were incubated in 2% aqueous uranyl acetate at 4°C overnight. After rinsing twice in water, samples were dehydrated in increasing concentrations of ethanol,(30%,50%,70%,90%,100%) and a final dehydration step in 100% acetone (twice, 10 min each). Samples were infiltrated with 50% acetone 50% Epon overnight in a tight closed container (to avoid humidity). Finally they were infiltrated twice with pure Epon (1h each at room temperature) and then placed in molds with fresh Epon (EMBed 812, Electron Microscopy Sciences Cat 14 120). Blocks were heated at 56°C for 48 h. Semi-thin (0.5µm thick) and ultrathin sections (70 nm thick) were cut with a Leica UC7 ultramicrotome (Leica, Leica Microsystemes SAS, Nanterre, France). Semi-thin sections were stained with 1% toluidine blue in 1% borax and the region of interest was selected under the microscope. Ultra-thin sections of the selected region were collected on copper grids (200 mesh, EMS, Souffelweyersheim, France) and contrasted with Reynold’s lead citrate. ^50^ Ultrathin sections were observed with a Hitachi HT7700 electron microscope (Milexia, Saint Aubin 91190, France) operating at 100 kV. Pictures (4096x3008 pixels) were taken with a CMOS TEM camera microscope, NanoSprint12 (Milexia, Saint Aubin 91190, France). Pictures were processed with the open source image processing program ImageJ, NIH, Bethesda, USA, when needed.

### Dural cells isolation for scRNA-seq. analysis

Mice were euthanized and perfused with 20 mL of PBS. Skullcaps were dissected using small surgical scissors and immediately placed in ice-cold PBS. Dural meninges were carefully peeled off the skullcaps using fine forceps and transferred to tubes for enzymatic digestion in RPMI medium (Gibco, 21875-034) supplemented with 2.5 mg/mL collagenase D (Roche, 11088866001) and 0.1 mg/mL DNase I (Roche, 11284932001) for 20 minutes at 37°C. The tissue was then dissociated by repeated pipetting (25 strokes) and passed through a 35-μm cell strainer. Following centrifugation (5 minutes at 400g, 4°C), cells were resuspended in 150 μL PBS and pooled from two meninges per well in 96-well V-bottom plates.

### Flow cytometry for scRNA-seq. analysis

To minimize nonspecific antibody binding, cell preparations were first incubated for 10 minutes at 4°C with anti-CD16/32 (Fc block; BD Biosciences, 5 µg/mL). Subsequently, a fluorescently conjugated antibody cocktail, including Live-or-Dye 405/452 (Biotium, 32003, 0.05 µL/test), was added. Cells were incubated with the antibodies for 30 minutes at 4°C, followed by centrifugation (5 minutes at 500g, 4°C) and a single PBS wash. Finally, cells were resuspended in 600 µL of PBS and sorted using the MoFlo Astrios cell sorter (Beckman Coulter). Dural cells were identified as viable singlets and collected in PBS containing 20% BSA.

### scRNA-seq. analysis

singleCell library prep were realized with chromium from 10X Genomics. Chromium Next GEM Single Cell 3’ GEM kit and Chip G were used, following manufacturer’s recommendations. 20000 cells were loaded on the chromium and sequenced on ILLUMINA NovaseqXplus. Using 100 cycles cartridge from ILLUMINA, calculations are based on 80000 reads/cell minimum per sample. The datasets from Sham and JVL-14 mice were merged and processed together to perform clustering and downstream analyses. Cells were grouped into clusters based on similarities in gene expression profiles using principal component analysis (PCA) for dimensionality reduction. A range of resolutions (0–1.5, step 0.1) was tested to evaluate cluster stability, and a resolution of 0.6 was selected, resulting in 24 clusters. Uniform Manifold Approximation and Projection (UMAP) was used for visualizing clusters. Cluster-specific violin plots were generated to assess key metrics such as the number of genes detected (nFeature_RNA), RNA counts (nCount_RNA), and the percentage of mitochondrial and ribosomal genes. Clusters were annotated by comparing cluster-specific gene expression profiles to marker gene lists and reference datasets^51–54^. Preliminary annotations were verified and refined using the Quby platform. Differential expression of genes between conditions within clusters were performed on Quby using DESeq2. Gene Set Enrichment Analysis (GSEA) was conducted using the differentially expressed data using Quby. Pathways with a p value less than 0.05 were plotted. Receptor-ligand interaction analysis was performed with CellChat, an R toolkit designed for the quantitative inference and visualization of intercellular communication networks from scRNA-seq data. All fibroblast clusters and immune cell clusters with at least 50 cells in both conditions were selected for ligand-receptor analysis. Data visualization was done on R version 4.3.1 using the packages ggplot2, ComplexHeatmap, and the inbuilt visualization tools in CellChat. Interactions with significant differences in communication probability between conditions were selected.

### Statistical analyses

Statistical analyses were performed using GraphPad Prism (version 10.3.1, GraphPad Software, San Diego, CA, USA). The normality of distributions was evaluated using the Shapiro-Wilk and Levene’s tests. Data are presented as mean (SD), with error bars representing SD. For normally distributed data, unpaired t-tests, two-way ANOVA, or one-way ANOVA followed by Dunnett’s or Tukey’s multiple comparisons were used. For data that did not meet normality assumptions, Mann-Whitney U-tests or Kruskal-Wallis tests with Dunn’s multiple comparisons were applied. Statistical significance was set at p < 0.05.

## Acknowledgments

We thank Phil Colsh (Yale Neurology) for comments on the manuscript.

This work was supported by Venolymphatic (BBT.1300), NEIMO (FRN, Fond de Recherche Neurosciences), BrainWash (ANR-17-CE14-0005), Lymbrain (ANR-20-CE16-0027-01). The authors are funded by the following sources: M.-R. El Kamouh, Lymbrain; M. Spajer: BBT.1300 Venolymphatic; R. Singabahu: BBT.1300 Venolymphatic, INCA (Institut National du Cancer); Anne-Laure Joly Marolany: NEIMO (FRN); S. Lehericy, Investissements d’Avenir, IAIHU-06 (Paris Institute of Neurosciences—Institut Hospitalo Universitaire), ANR-11-INBS-0006; S. Lenck: Institut National de la Santé et de la Recherche Médicale and Assistance Publique des Hôpitaux de Paris CIHU-2021. All animal work was conducted at the PHENO-ICMice facility. The Core is supported by two “Investissements d’Avenir” grants (ANR-10-IAIHU-06 and ANR-11-INBS-0011-NeurATRIS) and the Fondation pour la Recherche Médicale. We are greatly indebted to the Institut du Cerveau et de la Moelle épinière -Quant (Confocal imaging), - Histomics (Histology), – iGenSeq (Genotyping and sequencing), – DAC (Data analysis core facility), -Vectorology, -ICV-3C (Culture Cellulaire & Cytométrie) and -CENIR (Brain imaging) platforms for their technical assistance.

## Author contributions

Study concept: S Lenck, J-L Thomas. Study coordination and manuscript writing: S Lenck, J-L Thomas and MR El Kamouh. Provision of reagents: J-L Thomas. Methodological advisors: A Eichman, M Mazighi, PM Lledo, H Benveniste, M Santin, S Lehéricy, S Lenck, JL Thomas. Interpretation of data: S Lenck, JL Thomas, MR El Kamouh, H Benveniste, KA Sailor, A Eichmann. Figures: S Lenck, MR El Kamouh, R Singhabahu, JL Thomas. Human MRI protocol: S Lenck, S Lehericy. Patient recruitment: S Lenck, A Grine. Human MRI analysis: S Lenck, D Doukhi, R. Singhabahu, A Nasry, E Buscher. Mice model of JVL: S. Lenck, MC Bourriene. Mice surgeries: M Spajer, S Lenck, MC Bourriene, AL Joly Marolany. Mice 11.7T MRI: M Santin, L Mouton, A Ruze, S Lenck, M Spajer. Mice DCE-MRI: H Benveniste, S Koundal, M Spajer. Mice MRI analysis: R Singhabahu, J Gottschalk, H Benveniste, S Koundal, S Lenck. Two-photons imaging: KA Sailor, J Ninneman, PM Lledo. Immunolabeling and confocal imaging: MR El Kamouh, R Singhabahu, D Akbar, AL Joly Marolany. Electronic Microscopy: D. Langui. LSFM: M. Spajer, R Singhabahu, AL Joly Marolany. RNA-seq preparation: MR El Kamouh, J Van Wassenhove. RNA dataset analysis: R Singhabahu, MR El Kamouh, JL Thomas. Tracer uptake: MR El Kamouh. Blood sampling and Bioplex analysis: MR El Kamouh, J Van Wassenhove. Data analyses and statistics: S Lenck, R Singhabahu, MR El Kamouh.

