## Supplementary figures for "Cerebral Venous Blood Flow Regulates Brain Fluid Clearance via Dural Lymphatics"

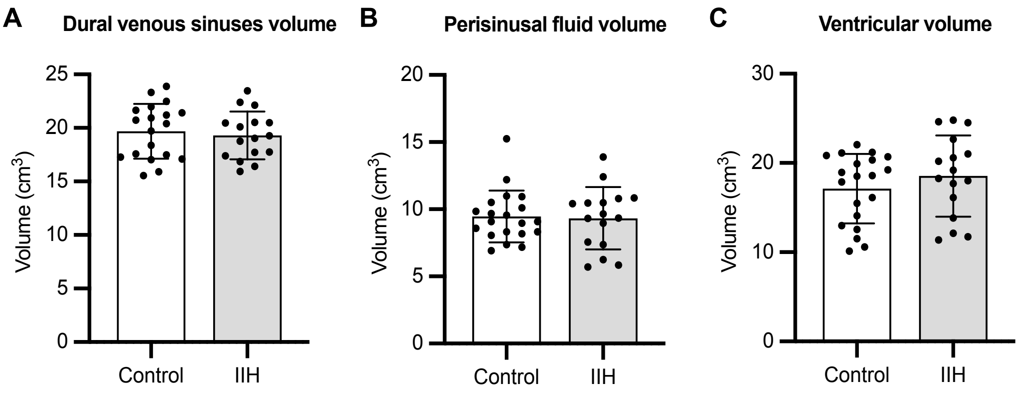


**SF1. MRI volumetric measurement in IIH patients and healthy controls.**
(A-C) Volume of dural venous sinuses (A), perisinusal space (B) and ventricules (C). Unpaired t-tests showed no significant difference between groups.

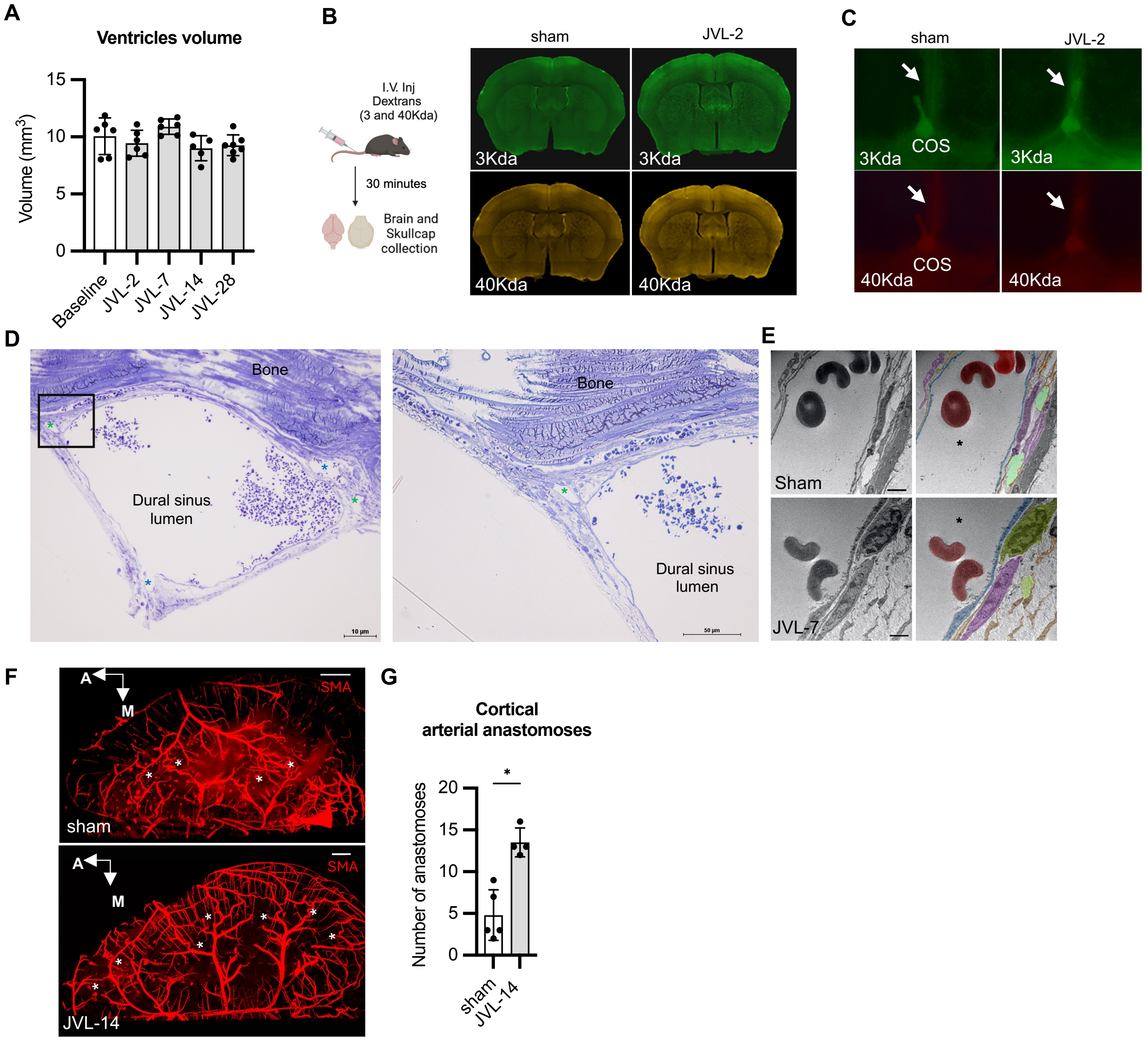


**SF2. Assessment of ventricular volume, BBB permeability, dural and brain vasculature in JVL mice.** (A) A 11.7 Tesla MRI scanner was use with a 3D-MGE sequence to measure ventricular volume, before and after JVL. (B, C) JVL-2 mice were tested for BBB integrity at one hour after IV. injection of 3- and 40-kDa dextrans. Extravasation of dextrans was detected neither in the brain parenchyma (B) nor in the dural perisinusal spaces (C). Superior sagittal sinus (white arrow). COS: Convergence of sinuses. (D) Electron microscopy of semi-thin sections from sham-operated mice showing the circulating lumen (Circ. Lumen) surrounded by the perisinus, with cortical vein entries (blue asterisks) and dural lymphatics (green asterisks) visible. (E) Ultrathin sections imaged by electron microscopy in sham-operated and JVL-7 mice reveal the microstructure of the dural sinus and confirm the integrity of the venous endothelium after JVL (blue). The circulating lumen (black asterisks) is bordered by a layer of venous endothelial cells (blue) and surrounded by the perisinus, composed of fibroblasts (pink), their extensions (orange) trapping collagen bundles (light green). No gaps are observed between the dural sinus lumen and the perisinus in JVL-7 mice. A macrophage (dark green) is noted in JVL-7 mice. Statistical analysis: ordinary one-way ANOVA with Dunnett's multiple comparisons test (A). (F,G) LSFM imaging of the cerebral arterial vasculature. Images of whole brain arterial tree after immunolabeling with smooth muscle actin (SMA) in sham operated and JVL-14 mice (F). Note the increased number of leptomeningeal anastomoses (stars) connecting branches of the middle and the anterior cerebral arteries in JVL mice. Corresponding quantification in (G). Mann Whitney U-test was used in G. * P<0.05. Elements in (B) were created with BioRender (https://biorender.com). Scale bars: 10 µm (D, left), 50 µm (D, middle), 5 µm (D, right), 1000 µm (F)

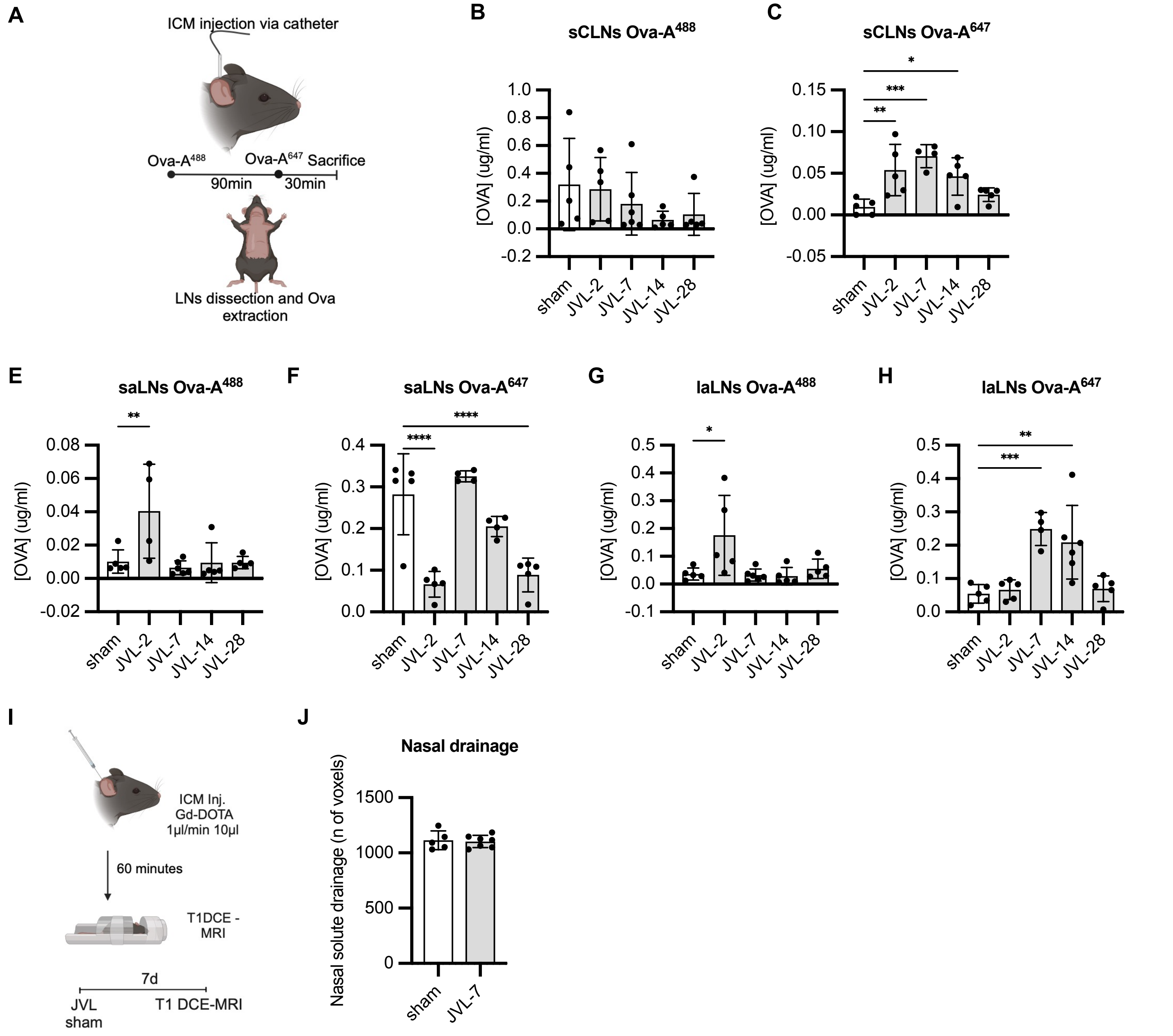


**SF3. Kinetics of CSF lymphatic drainage after jugular ligation.**

(A) CSF drainage into cervical and lumbo-sacral lymph nodes (LNs) was evaluated after ICM. injection of OVA-A$^{488}$ and OVA-A$^{647}$ at 120 min and 30 min, respectively, before sacrifice of JVL (-2, -7, -14 and -28) or sham-operated mice. (B-H). Fluctuation of fluorescent tracers collected in the superficial cervical lymph nodes (sCLNs) (B,C), the sacral lymph nodes (saLNs) (E,F) and the lumbo-aortic lymph nodes (laLNs) (G,H). (I, J) T1 DCE-MRI of the head of JVL-7 and sham-control mice at one hour after intracisternal injection of Gd-DOTA contrast agent. Schematic of the experiment (I). Quantitative analysis of Gd-DOTA signal across the cribriform plate showed no significant difference in CSF drainage between groups (n = 5 (sham), 7 (JVL)) (J). Ordinary one-way ANOVA with Dunnett's multiple comparisons test (B-H), Unpaired t-test (J). Elements in A and I were created using BioRender (https://biorender.com).

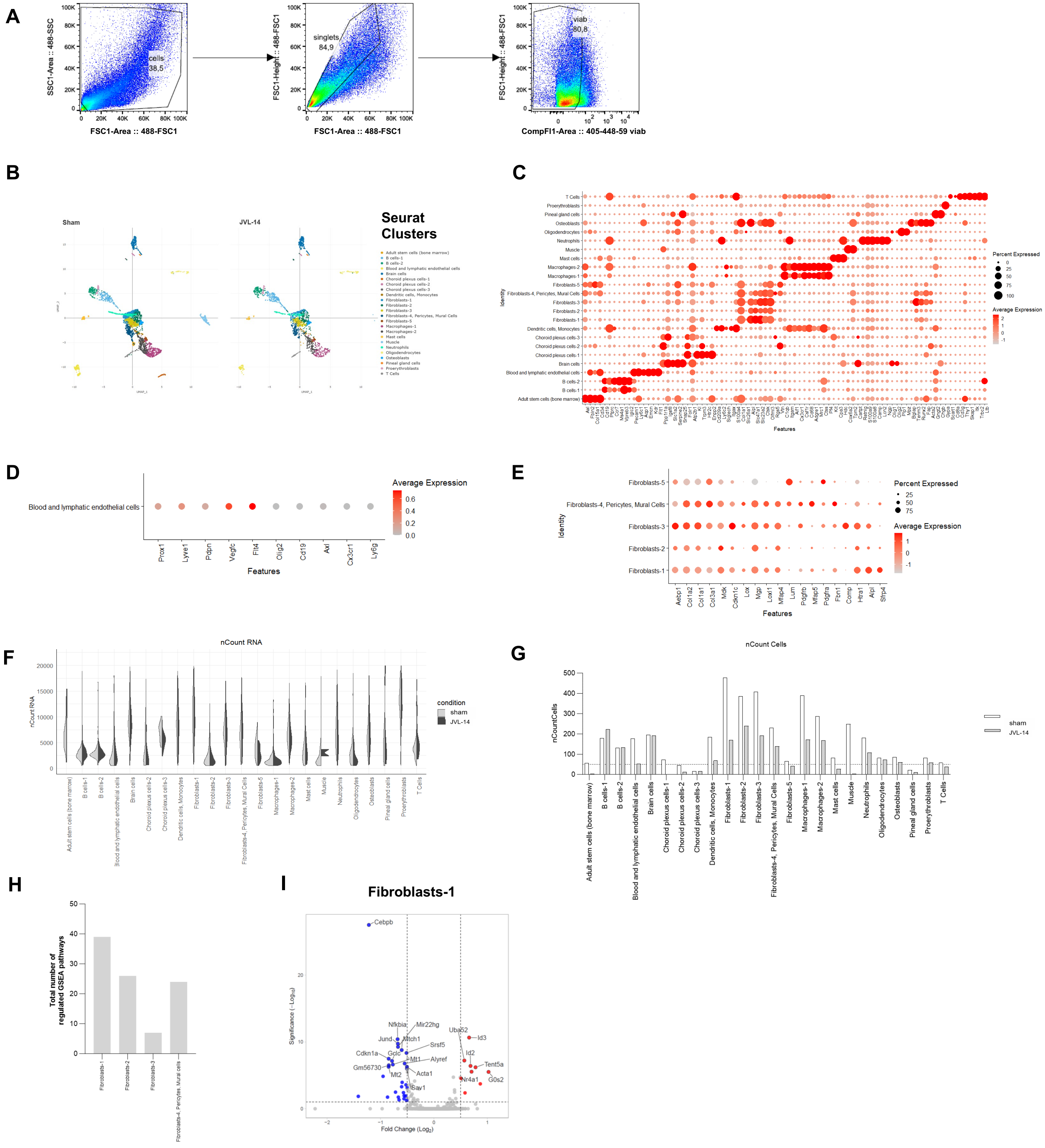


**SF4. FACS and scRNA-seq analysis of dura mater cells from sham-operated and JVL-14 mice.** (A) Flow cytometry gating strategy for sorting viable singlets. (B) Split 2D UMAP showing scRNA-seq identified dura mater clusters. (C) Dot plot of gene expression markers for each identified cell cluster. (D) Dot plot of selected genes including lymphatic markers within the Blood and lymphatic endothelial cells cluster. (E) Dot plot of selected genes within all fibroblast clusters. (F) Violin plots representing the number of transcripts by cluster. (G) Histogram of the cell counts among clusters. Dashed line represents the minimum threshold of 50 cells used for further GO enrichment analysis. (H) Histogram of the total number of regulated GSEA pathways in fibroblast clusters from JVL-14 and sham-operated mice. (I) Volcano-plot of up- and down-regulated genes in the Fibroblasts -1 cluster.
